# Calsyntenin-3, an atypical cadherin, suppresses inhibitory basket- and stellate-cell synapses but boosts excitatory parallel-fiber synapses in cerebellum

**DOI:** 10.1101/2021.05.31.446373

**Authors:** Zhihui Liu, Man Jiang, Kif Liakath-Ali, Jaewon Ko, Roger Shen Zhang, Thomas C. Südhof

## Abstract

Cadherins contribute to the organization of nearly all tissues, but the functions of several evolutionarily conserved cadherins, including those of calsyntenins, remain enigmatic. Puzzlingly, two distinct, non-overlapping functions for calsyntenins were proposed: As postsynaptic neurexin ligands in synapse formation, or as presynaptic adaptors for kinesin-mediated vesicular transport. Here, we show that acute CRISPR-mediated deletion of calsyntenin-3 in cerebellar Purkinje cells *in vivo* causes a large decrease in inhibitory synapses, but a surprisingly robust increase in excitatory parallel-fiber synapses. No changes in the dendritic architecture of Purkinje cells or in climbing-fiber synapses were detected. Thus, by promoting formation of an excitatory type of synapses and decreasing formation of an inhibitory type of synapses in the same neuron, calsyntenin-3 functions as a postsynaptic adhesion molecule that regulates the excitatory/inhibitory balance in Purkinje cells. No similarly opposing function of a synaptic adhesion molecule was previously observed, suggesting a new paradigm of synaptic regulation.

## INTRODUCTION

Synapses mediate information transfer between neurons in brain, and process the information during transfer. In processing information, synapses are dynamic: Synapses are not only continuously restructured by various forms of synaptic plasticity, but are also eliminated and newly formed throughout life (Attardo et al., 2015; Pfeiffer et al., 2018). Synapse formation, elimination, and remodeling are thought to be organized by synaptic adhesion molecules (SAMs) (Südhof, 2021). Many candidate SAMs have been described, but most SAMs appear to make only partial contributions to the formation and specification of synapses. In particular, few SAMs were consistently found to contribute to the initial formation of synapses. At present, only adhesion-GPCR SAMs, such as latrophilins and BAIs, are known to have a major impact on synapse numbers when tested using rigorous genetic approaches (Anderson et al., 2017; Bolliger et al., 2011; Kakegawa et al., 2015; Sando et al., 2019; Sando and Sudhof, 2021; Sigoillot et al., 2015; Wang et al., 2020). In contrast, the majority of well-characterized SAMs, most prominently neurexins and LAR-type receptor phosphotyrosine phosphatases, appear to perform no major roles in establishing synaptic connections. Instead, these SAMs are essential for conferring onto synapses specific properties that differ between various types of synapses in a neural circuit (Chen et al., 2017; Emperador-Melero et al., 2021; Fukai and Yoshida, 2020; Missler et al., 2003; Sclip and Sudhof, 2020).

Calsyntenins (a.k.a. alcadeins) are atypical cadherins that are encoded by three genes in mammals (*Clstn1*-*3* in mice) and a single gene in Drosophila, C. elegans, and other invertebrates (Araki et al., 2003; Hintsch et al., 2002; Ohno et al., 2014; Vogt et al., 2001). Calsyntenins are type I membrane proteins containing two N-terminal cadherin domains followed by a single LNS-domain (also referred to as LG-domain), a transmembrane region, and a short cytoplasmic tail. Calsyntenins are primarily expressed in neurons, although a calsyntenin-3 (Clstn3) variant with a different non-cadherin extracellular domain is present in adipocytes (referred to as Clstn3β; Zeng et al., 2019).

Two different views of calsyntenin functions have emerged. The first view posits that calsyntenins are postsynaptic adhesion molecules that bind to presynaptic neurexins to mediate both excitatory and inhibitory synapse formation, whereas the second view proposes that calsyntenins are presynaptic adaptor proteins that mediate kinesin function in axonal transport. Extensive evidence supports both views.

In support of a role for calsynteins as a postsynaptic adhesion molecule, all calsyntenins were localized by immunoelectron microscopy to the postsynaptic densities of excitatory synapses in the cortex and cerebellum (Hintsch et al., 2002; Vogt et al., 2001). Moreover, calsyntenins induce presynaptic specializations in heterologous synapse formation assays when expressed in non-neuronal cells (Pettem et al., 2013). Most importantly, knockout (KO) mice of all three calsyntenins exhibit synaptic impairments (Kim et al., 2020; Lipina et al., 2016; Pettem et al., 2013; Ster et al., 2014). Careful analyses revealed that *Clstn3* KO mice exhibit a 20-30% decrease in excitatory synapse density in the CA1 region of the hippocampus (Kim et al., 2020; Pettem et al., 2013). In addition, Pettem et al. (2013) observed a similar decrease in inhibitory synapse density, although Kim et al. (2020) failed to detect the same decrease. Moreover, Pettem et al. (2013) detected a 30-40% decrease in mEPSC and mIPSC frequency, but unexpectedly found no change in excitatory synaptic strength as measured by input/output curves. The *Clstn2* KO also decreased the inhibitory synapse density in the hippocampus approximately 10-20% (Lipina et al., 2016), whereas the *Clstn1* KO modestly impaired excitatory synapses in juvenile but not in adult mice (Ster et al., 2014). Viewed together, these data suggest a postsynaptic role for calsyntenins in the hippocampus, although the modest effect sizes of the calsyntenin KO phenotypes were puzzling. Calsyntenins were proposed to function as postsynaptic adhesion molecules by binding to presynaptic neurexins, but distinct, mutually exclusive mechanisms of neurexin binding were described. Pettem et al. (2013) and Lu et al. (2014) showed that the LNS domain of calsyntenins binds to an N-terminal sequence of α-neurexins that is not shared by β-neurexins. Kim et al. (2020), however, demonstrated that the cadherin domains of calsyntenins bind to the 6^th^ LNS domain of neurexins that is shared by α- and β-neurexins. Adding to this puzzle, Um et al. (2014) did not detect direct binding of calsyntenins to neurexins, and no study has reconstituted a stable calsyntenin-neurexin complex.

Similar to the hypothesis that calsyntenins act as postsynaptic adhesion molecules, the proposal that calsyntenins function as presynaptic kinesin-adaptor proteins that facilitate vesicular transport is also based on extensive studies. This proposal focuses on the transport of vesicles containing APP (Araki et al., 2007; Konecna et al., 2006; Vagnoni et al., 2012). A cytoplasmic sequence of calsyntenins binds to kinesins (Konecna et al., 2006), and at least *Clstn1* is localized to transport vesicles containing kinesin (Ludwig et al., 2009). Moreover, carefully controlled immunoprecipitations showed that calsyntenins are present in a molecular complex with presynaptic GABA_B_-receptors and APP (Dinamarca et al., 2019; Schwenk et al., 2016). However, *Clstn1* KO mice exhibit only modest changes in APP transport or the proteolytic processing of APP into Aβ peptides, while *Clstn2* KO mice display no change (Gotoh et al., 2020). Furthermore, the *Clstn1* KO simultaneously increased the levels of the C-terminal cleavage fragment (CTF) of APP and of Aβ peptides without changing APP levels, making it difficult to understand how a decreased APP cleavage causing increased CTF levels could also elevate Aβ levels (Gotoh et al., 2020). As a result, the kinesin binding, APP interaction, and GABA_B_-receptor complex formation by calsyntenins seem well established, but it is not yet clear how these activities converge on a function in axonal transport of APP.

The two divergent views of the function of calsyntenins, although well supported, are difficult to reconcile with each other. Given the potential importance of calsyntenins, we here pursued an alternative approach to study their functions. We aimed to identify neurons that express predominantly one calsyntenin isoform in order to avoid potential redundancy, and then examined the function of that calsyntenin isoform using acute genetic ablations and synapse-specific electrophysiological analyses. Our results reveal that cerebellar Purkinje cells express only *Clstn3* at high levels. Using CRISPR/Cas9-mediated deletions, we unexpectedly found that the *Clstn3* KO in Purkinje cells upregulated excitatory parallel-fiber synapses and had no effect on excitatory climbing-fiber synapses, but suppressed inhibitory basket- and stellate-cell synapses. These results indicate that a particular synaptic adhesion molecule can support formation of one class of synapses but suppress formation of another class of synapses in the same neuron.

## RESULTS

### *Clstn3* is the predominant calsyntenin isoform in cerebellar Purkinje cells

To analyze physiologically relevant functions of calsyntenins, we aimed to identify a type of neuron that expresses a particular calsyntenin isoform at much higher levels than others. This was necessary to avoid functional redundancy among multiple calsyntenins. Since most previous studies on calsyntenin functions were performed in the hippocampus (Kim et al., 2020; Lipina et al., 2016; Pettem et al., 2013; Ster et al., 2014), we examined calsyntenin expression in the hippocampus using single-molecule *in situ* hybridizations. All three calsyntenins were expressed in the CA1 and CA3 regions and the dentate gyrus. In the CA1 region, *Clstn1* and *Clstn3* levels were highest, in the CA3 region all three calsyntenins were similarly abundant, and in the dentate gyrus, *Clstn1* and *Clstn2* were most strongly present (Figure 1A). These results suggest that most hippocampal neurons co-express multiple calsyntenin isoforms, which may account for the modest phenotypes observed with *Clstn1*, *Clstn2*, and *Clstn3* KO mice (Kim et al., 2020; Lipina et al., 2016; Pettem et al., 2013; Ster et al., 2014).

**Figure 1:**
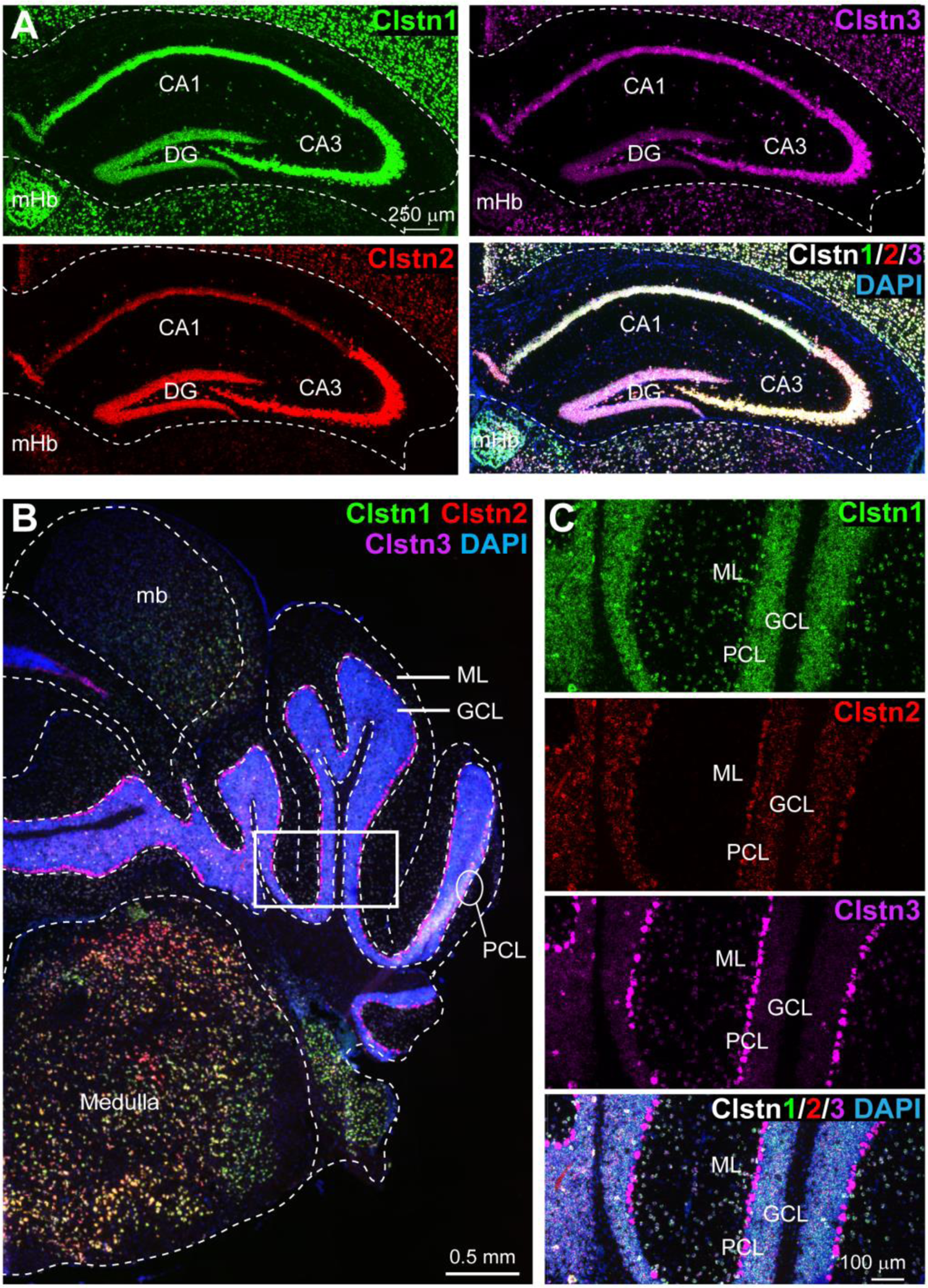
*Clstn1*, *Clstn2*, and *Clstn3* expression targets overlapping neuronal populations in the dorsal hippocampus, but distinct neuronal populations in the cerebellum of mice. **(A)** *Clstn1* (green), *Clstn2* (red) and *Clstn3* (magenta) exhibit distinct but largely overlapping expression patterns in the dorsal hippocampus. Representative images show sections from a mouse at P30 labeled by single-molecule *in situ* fluorescent hybridization (RNAscope) and by DAPI staining as indicated (DG, dentate gyrus; CA1 and CA3, CA1- and CA3-regions of the hippocampus proper; mHb, medial habenula). **(B & C)** *Clstn1* (green), *Clstn2* (red) and *Clstn3* (magenta) are expressed in separate and largely non-overlapping patterns in the cerebellum as visualized by single-molecule *in situ* hybridization (B, overview; C, expanded views of the area boxed in B; Mb, midbrain; ML, molecular layer; PCL, Purkinje cell layer; GCL, granule cell layer). Scale bars apply to all images in a set.

We next examined the cerebellum because single-cell RNA transcriptome databases suggested that in the cerebellum, granule cells express primarily *Clstn1*, while Purkinje cells express *Clstn3* (Peng et al., 2019; Saunders et al., 2018; Schaum et al., 2018; Tasic et al., 2018; Zeisel et al., 2018). Indeed, single-molecule *in situ* hybridization in cerebellar sections demonstrated that the expression of calsyntenin isoforms was much more segregated in cerebellum than in the hippocampus (Figure 1B). Purkinje cells express almost exclusively *Clstn3*, although low amounts of *Clstn2* were also detectable, whereas granule cells express *Clstn1* (Figure 1C). The differential labeling signal for the calsyntenins in the cerebellum was not due to differences in probe efficiency because in the hippocampus, the same probes under the same conditions produced equally strong signals (Figure 1A). Since Purkinje cells represent an excellent experimental system for synaptic physiology, we decided to focus on the function of *Clstn3* in these neurons.

### *In vivo* CRISPR efficiently deletes *Clstn3* in cerebellar Purkinje cells, causing major impairments in motor learning

Advances in CRISPR-mediated genetic manipulations suggest that it is possible to dissect a gene’s function using acute CRISPR-mediated deletions *in vivo* (Incontro et al., 2014). When we tested multiple single guide RNAs (sgRNAs) for the *Clstn3* gene, we identified two sgRNAs targeting exons 2 and 3 of the *Clstn3* gene that were highly efficacious in suppressing *Clstn3* expression in Purkinje cells *in vivo* (Figure 2A-2G). AAVs encoding the sgRNAs and tdTomato (as an expression marker) were stereotactically injected at P21 into the cerebellum of mice that constitutively express *sp*Cas9 (Platt et al., 2014), and mice were analyzed at ∼P50 by quantitative RT-PCR (Figure 2C, 2D). The *Clstn3* CRISPR KO was highly effective, decreasing *Clstn3* mRNA levels by >60% (Figure 2D, 2E). Immunoblotting confirmed the loss of ∼80% of *Clstn3* protein after the *Clstn3* CRISPR KO (Figure 2F). We also examined *Clstn1* and *Clstn2* expression by immunoblotting, using an antibody that reacts with both isoforms and additionally recognizes non-specific bands, but found no change (Figure S1). Analysis of the *Clstn3* CRISPR KO for potential off-target effects demonstrated that at the sites most similar to the *Clstn3* target sequence, no mutations were detected (Figure S2). Viewed together, these data indicate that the cerebellar *Clstn3* CRISPR KO effectively ablates *Clstn3* expression in cerebellar neurons.

**Figure 2:**
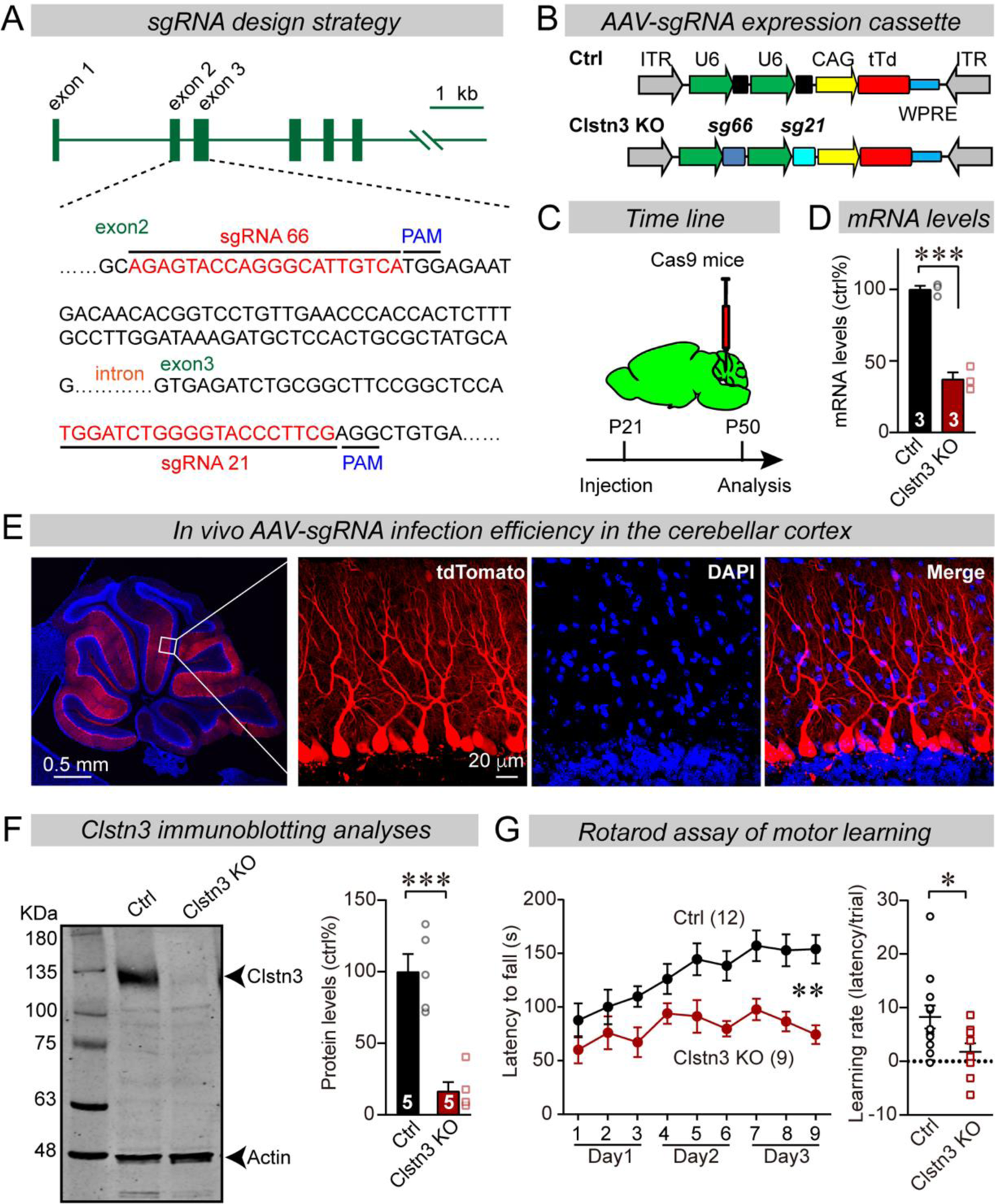
CRISPR/Cas9 manipulations enable rapid and highly efficient *in vivo* deletions of *Clstn3* in Purkinje cells, resulting in a severe impairment in motor learning. **(A)** Schematic of the sgRNA design strategy. Both sgRNAs target the positive strand of DNA, with sg66 targeting exon2, and sg21 targeting exon3. **(B)** Schematic of the AAV-DJ expression cassette in which sgRNAs and tdTomato (tdT) synthesis are driven by U6 and CAG promoters, respectively. Control mice were infected with AAVs that lacked sgRNAs but were otherwise identical. **(C)** Experimental strategy for CRISPR-mediated acute *Clstn3* deletions in the cerebellum. AAVs expressing the sgRNAs and tdTomato were stereotactically injected into the cerebellum of constitutively expressing Cas9 mice at P21, and mice were analyzed after P50. **(D)** Quantitative RT-PCR shows that the CRISPR-mediated *Clstn3* deletion severely suppresses *Clstn3* mRNA levels in the total cerebellum. Relative gene expression levels were first normalized to GAPDH using threshold cycle (CT) values, and then normalized to control. **(E)** Representative images of a cerebellar section from a mouse that was stereotactically infected with AAVs as described in **C** (left, overview of the cerebellum; right, cerebellar cortex; red = AAV-encoded tdTomato; blue, DAPI). Note that AAVs infect all Purkinje cells but few granule cells. **(F)** Immunoblotting analyses confirm that the CRISPR-mediated deletion greatly suppresses *Clstn3* protein levels (left, representative immunoblot; right, summary graph of quantifications using fluorescently labeled secondary antibodies). **(G)** The CRISPR-mediated *Clstn3* KO in cerebellar Purkinje cells severely impairs motor learning as analyzed by the rotarod assay (left, rotarod curve; right, slope of rotarod curve used as an index of the learning rate). Data in panels D, F, and G are means ± SEM. Statistical analyses were performed using unpaired t-test for D, F, and learning rate in G (*p<0.05, ***p<0.001) and repeat measures ANOVA for rotarod curve in G (F_(1,19)_=11.791, **p<0.01). Numbers of animals for each experiment are indicated in graphs.

To explore the functional consequences of the *Clstn3* KO in Purkinje cells, we analyzed its effect on the motor behavior of mice. Strikingly, the cerebellar *Clstn3* KO nearly abolished motor learning of the mice, as assayed with the rotarod task (Figure 2G). A behavioral test measuring social interactions did not reveal significant changes (Figure S3), suggesting that the mice were not broadly impaired. Thus, the *Clstn3* KO in the cerebellum likely impairs cerebellar functions in a specific manner. The motor coordination deficit in *Clstn3* CRISPR KO mice is consistent with phenotypes observed in constitutive *Clstn3* KO mice (www.mousephenotype.org) (Dickinson et al., 2016), suggesting that our CRISPR KO approach is reliable and specific.

### The *Clstn3* KO in Purkinje cells impairs inhibitory synapses in the cerebellar cortex

To test whether the *Clstn3* KO affected inhibitory synapses on Purkinje cells, we examined the inhibitory synapse density in the cerebellar cortex using immunocytochemistry for vGAT, an inhibitory synapse marker (Figure 3A). The CRISPR KO of *Clstn3* in Purkinje cells robustly reduced the inhibitory synapse density in the molecular layer, Purkinje-cell layer, and granule-cell layer of the cerebellar cortex (Figure 3B-3D). The most extensive decrease (∼60%) was observed in the deep molecular layer (Figure 3B), where we also detected a significant reduction (∼25%) in the size of vGAT-positive puncta (Figure 3C). vGAT-positive synapses in Purkinje cell and granule cell layers were less affected (∼20% reduction; Figure 3D).

**Figure 3:**
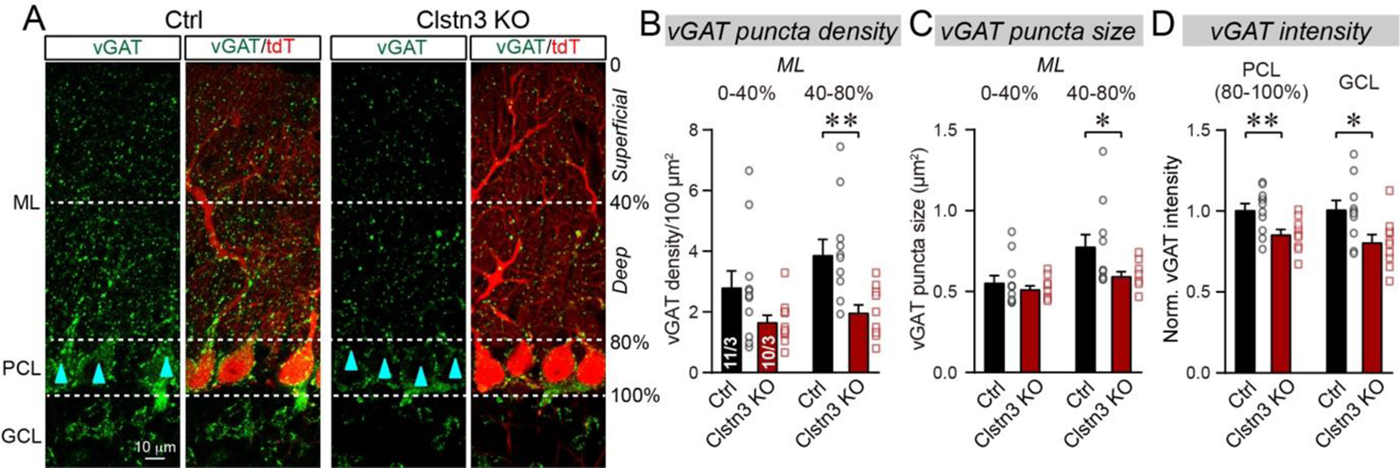
The *Clstn3* KO decreases inhibitory synapse numbers in the cerebellar cortex. **(A)** Representative confocal images of cerebellar cortex sections stained for vGAT and tdTomato. Sections are from mice in which Purkinje cells were infected with control AAVs (Ctrl) or AAVs that induce the CRISPR-mediated *Clstn3* KO (red, AAV-encoded tdTomato signal; green, vGAT; ML, molecular layer; PCL, Purkinje cell layer; GCL, granule cell layer). Calibration bar applies to all images. **(B-D)** The Clstn3 KO in Purkinje cells suppresses the number of vGAT-positive synapses in the cerebellar cortex. Summary graphs show quantifications of the density (B) and size (C) of vGAT-positive puncta in the molecular layer (ML) of the cerebellar cortex (separated into deep and superficial areas), and of the vGAT-staining intensity in the Purkinje cell layer (PCL) and granule cell layer (GCL) of the cerebellar cortex (D). Data are means ± SEM (numbers of sections/mice analyzed are indicated in bar graphs). Statistical analyses were performed using unpaired t-tests, with *p<0.05, **p<0.01.

The decrease in inhibitory synapse density raises the question whether inhibitory synaptic transmission is suppressed. To address this question, we recorded miniature inhibitory postsynaptic currents (mIPSCs) from Purkinje cells in the presence of tetrodotoxin (Figure 4). *Clstn3* KO produced a large decrease in mIPSC frequency (∼60%), without changing the mIPSC amplitude (Figure 4A-4C). Moreover, the *Clstn3* KO increased the rise but not decay times of mIPSCs (Figure 4D). Measurements of the Purkinje cell capacitance and input resistance showed that the Clstn3 deletion did not produce major changes, demonstrating that it did not globally alter Purkinje cell properties (Figure S4A).

**Figure 4:**
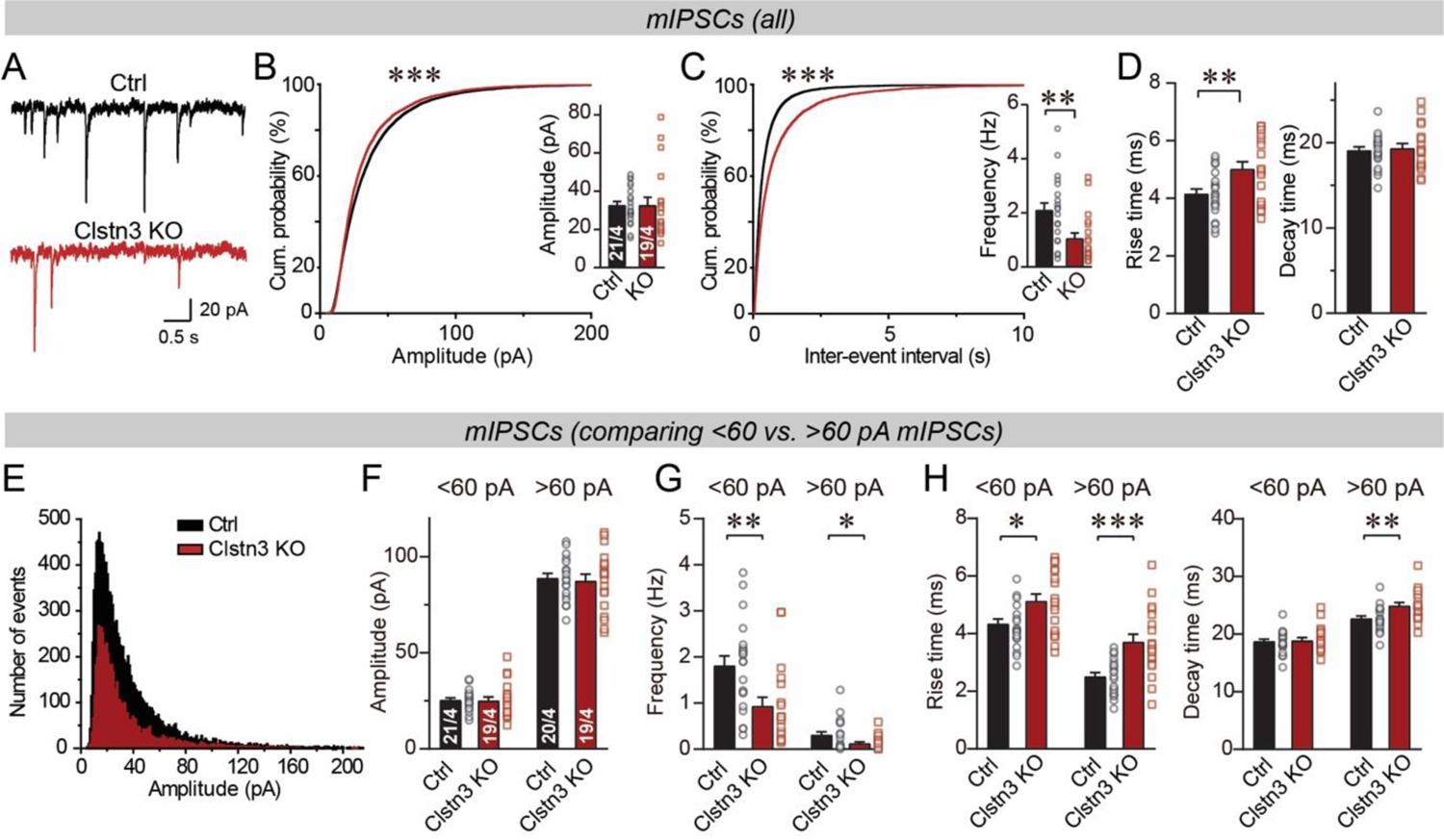
The *Clstn3* KO decreases spontaneous inhibitory synaptic ‘mini’ events in Purkinje cells. **(A-C)** The *Clstn3* KO decreases the frequency but not the amplitude of mIPSCs (A, representative traces; B, cumulative probability plot of the mIPSC amplitude [inset, average amplitude]; C, cumulative probability plot of the mIPSC inter-event interval [inset, average frequency]). **(D)** The Purkinje cell *Clstn3* KO increases the rise but not decay time of mIPSCs. **(E)** Plot of the number of mIPSC events vs. amplitude using a normal distribution. **(F-H)** The *Clstn3* KO similarly impairs mIPSCs with a larger (>60 pA) and a smaller amplitude (<60 pA), which in Purkinje cells are likely generated primarily by basket-cell and stellate-cell synapses, respectively (F & G, summary graphs for the mIPSC amplitude (F) and frequency (G) separately analyzed for high- and low-amplitude events; H, mIPSC rise [left] and decay times [right], separately analyzed for high- and low-amplitude events). All summary data are means ± SEM. Numbers of cells/mice analyzed are indicated in bar graphs. Statistical analyses were performed using unpaired t-tests (bar graphs with two groups) or Kolmogorov-Smirnov test (cumulative analysis), with *p<0.05, **p<0.01, ***p<0.001.

mIPSCs are heterogeneous in Purkinje cells. Smaller mIPSCs are mostly derived from more distant stellate-cell synapses, and larger mIPSCs from more proximal basket-cell synapses (Nakayama et al., 2012). To examine these two types of input synapses separately, we plotted the mIPSC amplitudes in a normal distribution (Figure 4E). This plot revealed that the majority of mIPSC amplitudes (>90%) are <60 pA. Therefore, we separately analyzed mIPSCs with amplitudes of >60 pA and <60 pA, of which the >60 pA mIPSCs likely represent basket cell mIPSCs, whereas the <60 pA mIPSCs are composed predominantly (but not exclusively) of stellate cell mIPSCs. Both classes of mIPSCs exhibited similar impairments in frequency and kinetics, although the changes were more pronounced for larger mIPSCs (Figure 4F-4H; S5). These results are consistent with the morphological data suggesting a larger decrease in basket cell synapses than in stellate cell synapses (Figure 3A-3D).

Does the change in inhibitory synapse density and mIPSCs cause a change in overall inhibitory synaptic strength? We examined evoked inhibitory synaptic responses, using extracellular stimulations of basket cell axons close to the Purkinje cell layer (Figure 5A). We detected a significant decrease (∼40%) in IPSC amplitudes. The decrease in IPSC amplitude is consistent with a loss of inhibitory synapses, but could also be due to a decrease in release probability. However, we detected no major changes in the coefficient of variation, paired-pulse ratio, or kinetics of evoked IPSCs, suggesting that the release probability is normal (Figure 5B-5F, S4B). These data confirm the morphological results, suggesting that the *Clstn3* KO decreases inhibitory synapse numbers.

**Figure 5:**
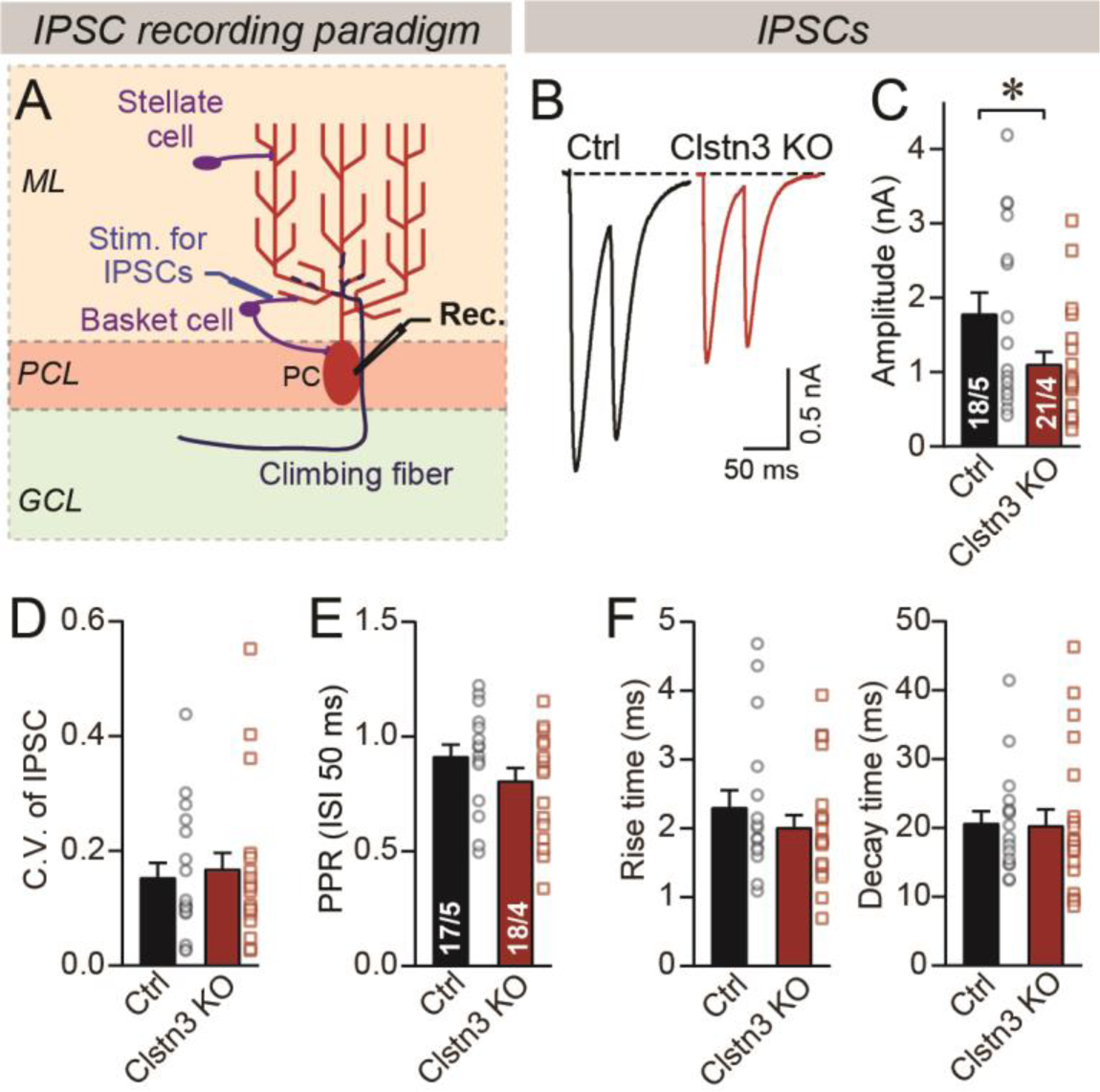
The *Clstn3* KO decreases evoked inhibitory synaptic responses in Purkinje cells. **(A)** Experimental design for recordings of IPSCs evoked by stimulation of basket cell axons (ML, molecular layer; PCL, Purkinje cell layer; GCL, granule cell layer; PC, Purkinje cell; Rec., recording patch pipette). **(B & C)** The *Clstn3* KO decreases the amplitude of evoked basket-cell IPSCs (B, representative traces of pairs of evoked IPSCs with a 50 ms inter-stimulus interval; C, summary graphs of the amplitude of the first IPSC). **(D & E)** The *Clstn3* KO in Purkinje cells does not affect the release probability at inhibitory synapses as judged by the coefficient of variation (D) and the paired-pulse ratio with an interstimulus interval of 50 ms (E) of evoked IPSCs. **(F)** The *Clstn3* KO in Purkinje cells has no significant effect on IPSC kinetics (left, rise times; right, decay times of evoked ISPCs). All summary data are means ± SEM. Numbers of cells/mice analyzed are indicated in bar graphs. Statistical analyses were performed using unpaired t-tests, with *p<0.05.

### *Clstn3* deletion in Purkinje cells increases excitatory parallel-fiber but not climbing-fiber synapse densities

The decrease in inhibitory synapse numbers by the *Clstn3* KO is consistent with previous studies suggesting that *Clstn3* promotes synapse formation in the hippocampus, but these previous studies primarily identified a decrease in excitatory synapses (Kim et al., 2020; Pettem et al., 2013; Ranneva et al., 2020). We thus tested whether the *Clstn3* KO also affects excitatory synapse numbers in cerebellum. Purkinje cells receive two different excitatory synaptic inputs with distinct properties: Parallel-fiber synapses that are formed by granule cells on distant Purkinje cell dendrites, and climbing-fiber synapses that are formed by inferior olive neurons on proximal Purkinje cell dendrites. Parallel-fiber synapses use the vesicular glutamate transporter vGluT1, whereas climbing-fiber synapses use the vesicular glutamate transporter vGluT2 (Hioki et al., 2003). Moreover, parallel-fiber synapses are surrounded by processes formed by Bergmann astroglial cells, creating a tripartite synapse in which the glial processes contain high levels of GluA1 (Baude et al., 1994). As a first step towards assessing the effect of the *Clstn3* KO on excitatory synapses on Purkinje cells, we analyzed cerebellar sections from control and *Clstn3* KO mice by immunohistochemistry for vGluT1, vGluT2 and GluA1 (Figure 6).

**Figure 6:**
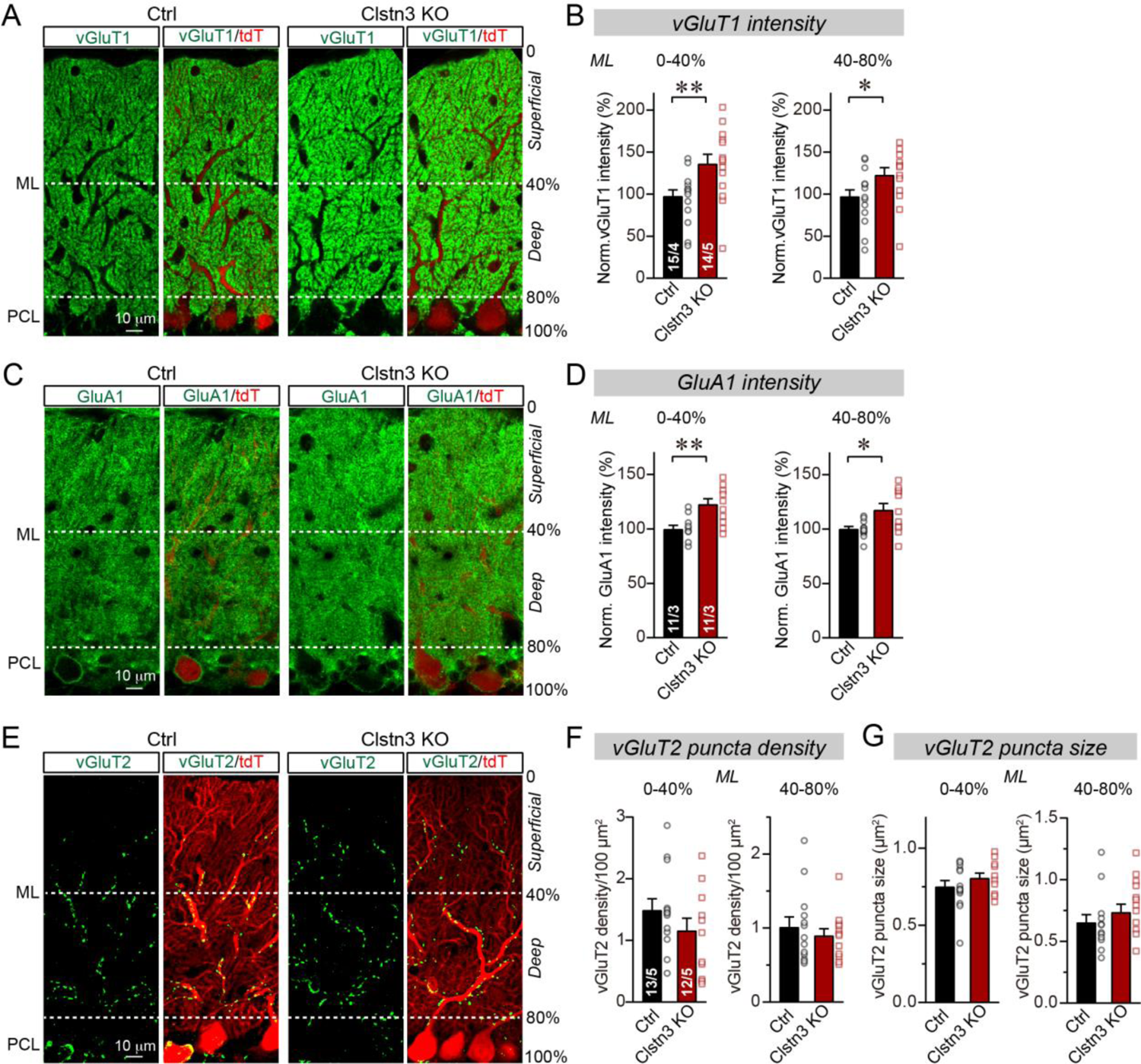
CRISPR-mediated *Clstn3* deletion in the cerebellar Purkinje cells increases parallel-fiber excitatory synapse numbers without changing climbing-fiber synapse numbers. **(A & B)** Immunostaining of cerebellar cortex sections with antibodies to vGluT1 as a presynaptic marker for parallel-fiber synapses reveals a significant increase (A, representative confocal images from control and *Clstn3* KO mice [green vGluT1; red, tdTomato]; B, summary graphs of the vGluT1 staining intensity in the superficial (0-40%) and deep (40-80%) molecular layers of the cerebellar cortex). (**C & D**) Immunostaining with antibodies to GluA1, an astroglial marker for tripartite parallel-fiber/Bergmann glia synapses, also uncovers a significant increase in staining intensity (C, representative confocal images from control and *Clstn3* KO samples [green vGluT1; red, tdTomato]; D, summary graphs of the GluA1 staining intensity in the superficial (0-40%) and deep (40-80%) molecular layers of the cerebellar cortex). **(E-G)** Immunostaining for vGluT2 as a marker for climbing-fiber synapses in cerebellar cortex fails to uncover a *Clstn3* KO-induced change (E, representative confocal images [green, vGluT2; red, tdTomato]; F & G, summary graphs of the density (F) and size (G) of vGluT2-positive synaptic puncta in the superficial (0-40%) and deep (40-80%) molecular layers of the cerebellar cortex). All numerical data are means ± SEM; numbers of sections/mice analyzed are indicated in the first bar graphs for each experiment. Statistical significance was assessed by unpaired Student’s t-test (*p<0.05, **p<0.01).

Confocal microscopy of cerebellar cortex sections immunolabeled for vGluT1 revealed intense staining that, surprisingly, was enhanced by the *Clstn3* deletion in Purkinje cells (Fig. 6A). Because parallel-fiber synapses in the cerebellar cortex are so numerous that confocal microscopy cannot resolve individual vGluT1-positive synaptic puncta, we measured the overall vGluT1 staining intensity as a proxy for synapse density (Figure 6A) (Zhang et al., 2015). The *Clstn3* deletion in Purkinje cells caused a robust increase (∼25%) in the vGluT1 staining intensity of both the superficial and the deep molecular layers of the cerebellar cortex (Figure 6B).

The potential increase in parallel-fiber synapses induced by the *Clstn3* KO, suggested by the enhanced vGluT1 staining intensity, is unexpected. This prompted us to examine the levels of GluA1 as an astroglial marker of tripartite parallel-fiber synapses (Figure 6C; Baude et al., 1994). Again, the *Clstn3* KO in Purkinje cells induced a significant increase (∼25%) in synaptic GluA1 staining intensity (Figure 6D), consistent with the increase in vGluT1 staining intensity.

We next analyzed the density of climbing-fiber synapses by staining cerebellar sections for vGluT2, but detected no significant effect of the *Clstn3* KO in Purkinje cells (Figure 6E). Different from parallel-fiber synapses that contain vGluT1, climbing-fiber synapses are labeled with antibodies to vGluT2 and are readily resolved by confocal microscopy (Figure 6E). The number and size of synaptic puncta identified with vGluT2 antibodies were not altered by the *Clstn3* KO, although there was a slight trend towards a decrease in climbing-fiber synapse density (Figure 6F, 6G). These observations suggest that the enhancement of parallel-fiber synapse density by the *Clstn3* KO is specific for this type of synapse.

### The *Clstn3* KO increases the spine density of Purkinje cells

It is surprising that the *Clstn3* KO in Purkinje cells appears to increase the parallel-fiber synapse density, as one would expect a synaptic adhesion molecule to promote but not to suppress formation of a particular synapse. The parallel-fiber synapse increase is likely not a homeostatic response to the loss of inhibitory synapses because such a response, which would aim to maintain the correct excitatory/inhibitory balance, should involve a decrease, and not an increase, in parallel-fiber synapses. The increase in parallel-fiber synapse numbers is also unexpected given previous results showing that in hippocampal CA1 neurons, the *Clstn3* KO decreases excitatory synapse numbers (Kim et al., 2020; Pettem et al., 2013). To independently confirm this increase, we analyzed the dendritic spine density in Purkinje cells. Since nearly all spines contain parallel-fiber synapses and all parallel-fiber synapses are on spines (Sotelo, 1975), the spine density of Purkinje cells represents a reliable proxy for synapse density.

We filled individual Purkinje cells in acute slices with biocytin via a patch-pipette, and analyzed their dendritic structure and spine density by quantitative morphometry (Figure 7A, S6A, S6B). Reconstructions of 6 Purkinje cells from control and *Clstn3* KO mice revealed a trend towards an increased dendrite length in *Clstn3*-deficient Purkinje cells without a significant change in dendritic architecture, demonstrating that the *Clstn3* KO does not impair the overall structure of Purkinje cells (Figure 7B). Quantification of dendritic spines uncovered in *Clstn3*-deficient Purkinje cells a robust increase (∼30%) in the density of spines in the superficial area of the cerebellar cortex, and a trend towards an increase in the deep area of the cerebellar cortex (Figure 7C-7F). The increase in spine density was particularly pronounced for thin spines (Figure 7G; S6C, S6D). These findings provide independent evidence that the *Clstn3* KO increases the parallel-fiber synapse density, and precisely mirror those obtained by analyzing the vGluT1- and GluA1-staining intensity of the cerebellar cortex (Figure 6A-6D).

**Figure 7:**
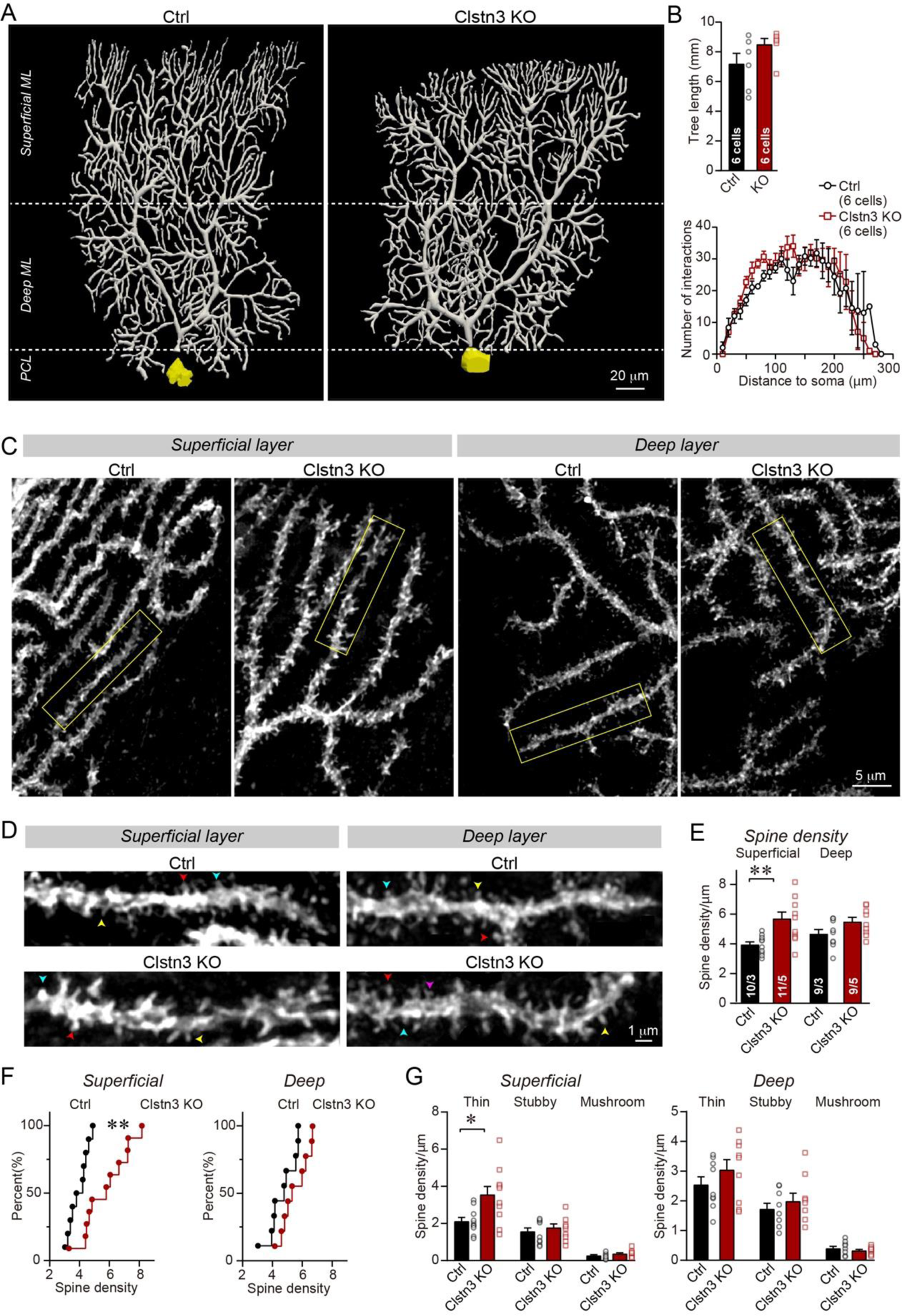
Morphological analysis of individual Purkinje cells reveals that the *Clstn3* KO robustly increases their dendritic spine density without significantly altering their dendritic arborization. **(A & B)** Biocytin filling of individual Purkinje cells via a patch pipette demonstrates that the *Clstn3* KO does not significantly change the overall dendritic architecture of Purkinje cells (A, representative images of Purkinje cell dendritic trees for control and *Clstn3* KO mice after 3D reconstruction [for more images, see Supplementary Fig. S6]; B, quantifications of the dendritic tree length [top] or dendritic arborization using Sholl analysis [bottom]). (**C-F**) The *Clstn3* KO increases the density of dendritic spines of Purkinje cells in the superficial part of the cerebellar cortex (C & D, representative images of spiny dendrites at low and high magnifications, respectively; [blue, red, and yellows arrowheads mark different spine types]; E & F, summary graph [E] and cumulative distribution of the spine density [F]). **(G)** The *Clstn3* KO in Purkinje cells increases preferentially the density of thin spines in the superficial part of the cerebellar cortex, based on a morphological classification of spine types into thin, stubby and mushroom spines. All data in B, E, and G are means ± SEM; 6 control and Clstn3 KO Purkinje cells were reconstructed for B; numbers in the first bars of E indicate the number of cell/animal analyzed for E-G. Statistical significance (*p<0.05; **p<0.01) in B and G was assessed by an unpaired t-test, and in E by one-way ANOVA (F_(3, 35)_=5.693, p=0.003), followed by Tukey’s post hoc comparisons for control and *Clstn3* KO groups.

### The *Clstn3* KO increases parallel-fiber but not climbing-fiber synaptic transmission

The increase in parallel-fiber synapses could be due to a true enhancement of parallel-fiber synapse formation, or a compensatory reaction to a decrease in parallel-fiber synapse function. Although the latter hypothesis would be consistent with a homeostatic response, it seems unlikely given that in vertebrates, synapses rarely proliferate in response to a functional impairment. To clarify this question, we analyzed parallel-synapse function by electrophysiology, and compared it to climbing-fiber synapse function as an internal control since climbing-fiber synapse numbers are not changed by the *Clstn3* KO in Purkinje cells.

We first monitored spontaneous miniature synaptic events (mEPSCs) in the presence of tetrodotoxin. We observed an increase in mEPSC amplitudes (∼25%) and frequency (∼15%) in *Clstn3* KO neurons, without a notable change in mEPSC kinetics (Figure 8A-8D). Most mEPSCs in Purkinje cells are derived from parallel-fiber synapses. Because of the large dendritic tree of Purkinje cells, synapses on distant dendrites produce slower and smaller mEPSCs than synapses on proximal dendrites (Zhang et al., 2015). To ensure that we were monitoring mEPSCs derived from parallel-fiber synapses (whose density is increased morphologically), we analyzed only slow mEPSCs with rise times of >1 ms that are mostly generated by parallel-fiber synapses on distant dendrites (Nakayama et al., 2012; Yamasaki et al., 2006). The results were the same as for total mEPSCs, confirming that the *Clstn3* KO increases parallel-fiber synaptic activity (Figure 8E-8H).

**Figure 8:**
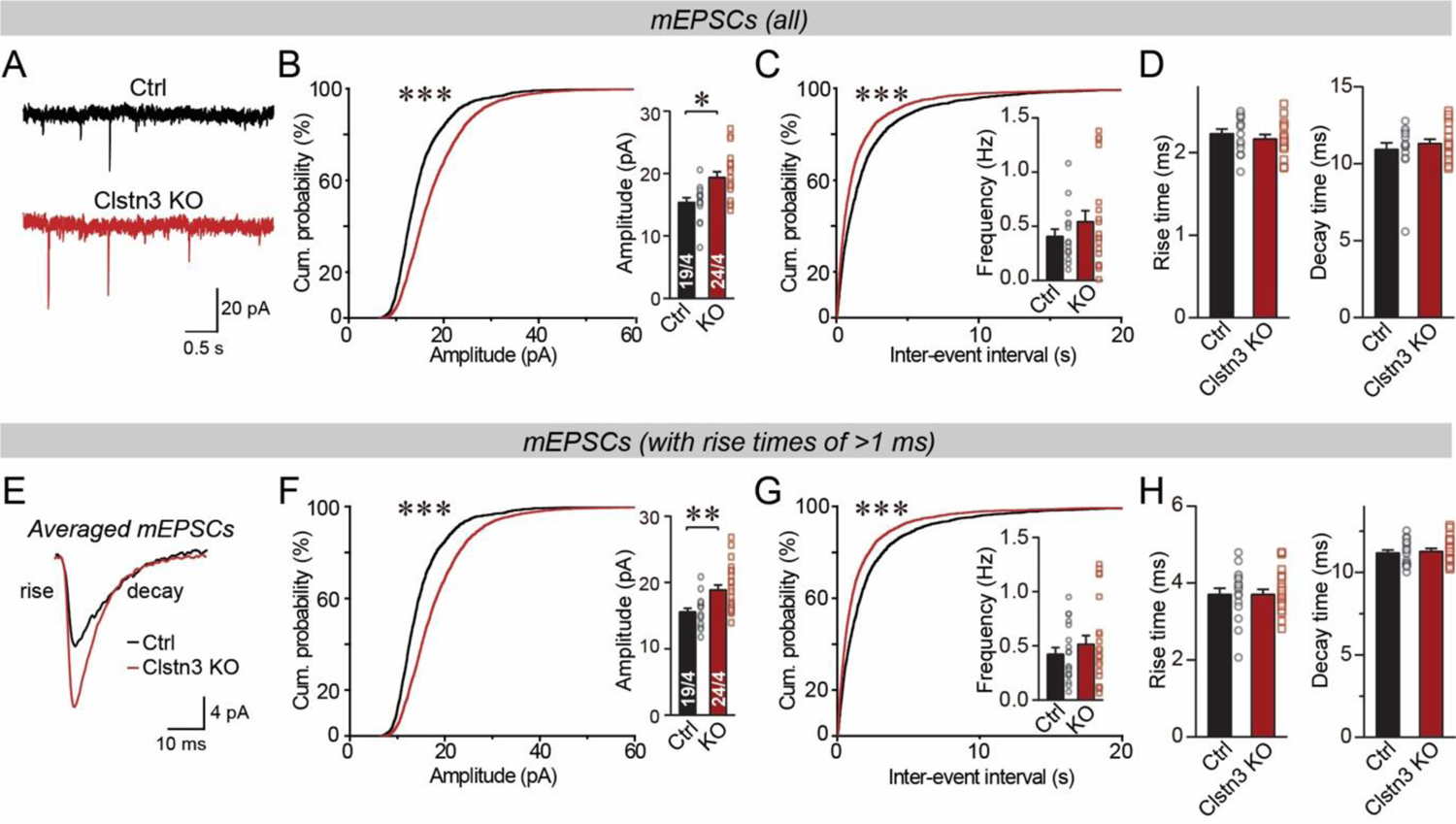
The *Clstn3* KO in cerebellar Purkinje cells increases the amplitude and frequency of parallel-fiber mEPSCs. **(A-C)** The *Clstn3* KO increases the amplitude and frequency of mEPSCs in Purkinje cells (A, representative traces; B, cumulative probability plot of the mEPSC amplitude [inset, average amplitude]; C, cumulative probability plot of the mEPSC inter-event interval [inset, average frequency]). **(D)** The *Clstn3* KO in Purkinje cells has no effect on mEPSC kinetics (left, mEPSC rise times; right, mEPSC decay times). **(E)** Expanded trace of averaged mEPSCs to illustrate the kinetic similarity of control and *Clstn3* KO events with a change in amplitude. **(F & G**) mEPSCs with slow rise times (>1 ms) and that are likely exclusively derived from parallel-fiber synapses exhibit the same phenotype as the total mEPSCs (same as B & C, but for mEPSCs with slow rise times). **(H)** The *Clstn3* KO in Purkinje cells has no effect on the kinetics of mEPSCs with slow rise times (left, mEPSC rise times; right, mEPSC decay times). All numerical data are means ± SEM. Statistical significance with two groups was assessed by unpaired t-test (*p<0.05, **p<0.01), with the number of cells/mice analyzed indicated in the first bar graphs for each experiment. Cumulative analysis was done with Kolmogorov-Smirnov test (***p<0.001).

Finally, we measured evoked parallel-fiber EPSCs, using input-output curves to correct for variations in the placement of the stimulating electrode (Figure 9A). Consistent with the morphological and mEPSC data, the *Clstn3* KO robustly enhanced parallel-fiber synaptic responses (∼60% increase) (Figure 9B-9D). This finding suggests that the *Clstn3* KO not only increases the density of parallel-fiber synapses, but also renders these synapses more efficacious. The increased strength of parallel-fiber synaptic transmission was not due to a change in release probability because neither the coefficient of variation nor the paired-pulse ratios of parallel-fiber EPSCs were affected (Figure 9E-9G). The increase of parallel-fiber EPSCs is consistent with the vGluT1 intensity and mEPSC amplitude changes, providing further evidence that the *Clstn3 KO* enhances parallel-fiber synapses.

**Figure 9:**
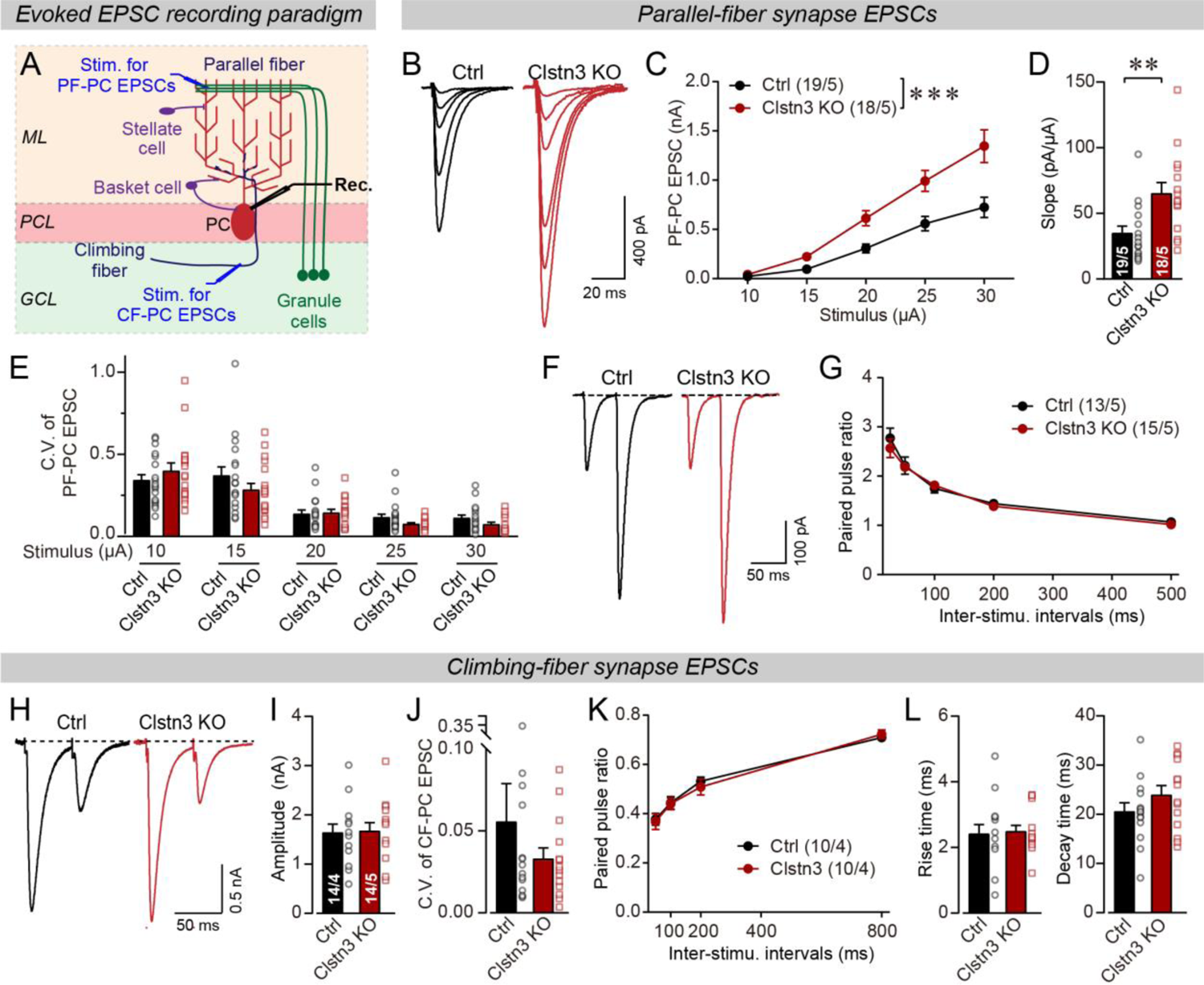
*Clstn3* KO in Purkinje cells elevates parallel-fiber synaptic strength, while leaving climbing-fiber synaptic strength unchanged. **(A)** Schematic of the recording configuration for monitoring evoked EPSCs induced by parallel-fiber (PF-PC) and climbing-fiber stimulation (CF-PC) in Purkinje cells. **(B-D)** The postsynaptic *Clstn3* KO robustly increases the input/output relation of parallel-fiber synapses (B, representative traces; C, input/output curve; D, summary graph of the slope of the input/output curve determined in individual cells). **(E-G)** The postsynaptic *Clstn3* KO in Purkinje cells has no effect on presynaptic release probability as assessed by monitoring the coefficient of variation of evoked EPSCs (E, separately analyzed for different stimulus strengths) or the paired-pulse ratio (F, sample traces; G, plot of the paired-pulse ratio of parallel-fiber EPSCs as a function of interstimulus interval). **(H-L)** The *Clstn3* KO has no effect on the amplitude, coefficient of variation, paired-pulse ratio, or kinetics of climbing-fiber synapse EPSCs (H, representative traces of climbing-fiber EPSCs elicited with an interstimulus interval of 50 ms; I & J, amplitude (I) and coefficient of variation (J) of evoked climbing-fiber EPSCs; K, plot of the paired-pulse ratio of climbing-fiber EPSCs as a function of interstimulus interval; L, rise [left] and decay times [right] of evoked climbing-fiber EPSCs). All numerical data are means ± SEM. Statistical analyses were performed by two-way ANOVA followed by Tukey’s post hoc correction (C, G, K; for C, F_(1, 150)_=35.83, p<0.0001) or unpaired t-test for experiments with two groups (D, E, I, J, L), with *p<0.05, **p<0.01.

In contrast to parallel-fiber EPSCs, climbing-fiber EPSCs exhibited no *Clstn3* KO-induced alteration. Specifically, the amplitude, paired-pulse ratio, and kinetics of climbing-fiber EPSCs in control and *Clstn3* KO Purkinje cells were indistinguishable (Figure 9H-9L). These findings are consistent with the lack of a change in vGluT2-positive synaptic puncta analyzed morphologically (Figure 6E-6G). Viewed together, these data suggest that *Clstn3 KO* produces an increase in excitatory parallel-fiber, but not climbing-fiber, synapses.

## DISCUSSION

Calsyntenins are intriguing but enigmatic cadherins, a class of diverse adhesion molecules that generally function as tissue organizers. Two distinct, non-overlapping roles were proposed for calsyntenins, as a postsynaptic adhesion molecule promoting synapse formation and as a kinesin-adaptor protein mediating axonal and dendritic transport, in particular of APP (Araki et al., 2007; Kim et al., 2020; Konecna et al., 2006; Lipina et al., 2016; Pettem et al., 2013; Ster et al., 2014; Vagnoni et al., 2012). Both functions are supported by extensive data, but neither function was conclusively tested. Here, we examined the role of one particular calsyntenin, *Clstn3*, in one particular neuron, Purkinje cells that predominantly express this calsyntenin isoform. Our data establish that *Clstn3* acts as a postsynaptic adhesion molecule in Purkinje cells that is selectively essential for regulating synapse numbers, confirming an essential function for *Clstn3* as a synaptic adhesion molecule. Our data are surprising in revealing that *Clstn3* functions not by universally promoting synapse formation, but by exerting opposite effects in different types of synapses. Specifically, our results demonstrate that deletion of *Clstn3* causes a decrease in inhibitory basket- and stellate-cell synapses on Purkinje cells, but an increase in excitatory parallel-fiber synapses (Figure S7). Thus, *Clstn3* doesn’t function simply as a synaptogenic adhesion molecule, but as a regulator of the balance of excitatory and inhibitory synaptic inputs on Purkinje cells.

The functions we describe here for *Clstn3* are different from those of previously identified synaptic adhesion molecules or synapse-organizing signals. Whereas presynaptic adhesion molecules generally act in both excitatory and inhibitory synapses, few postsynaptic adhesion molecules were found to function in both. In the rare instances in which an adhesion molecule was documented to mediate signaling in excitatory and inhibitory synapses, such as the case of *Nlgn3* (but not of other neuroligins), it acts to promote synaptic function in both (Chanda et al., 2017; Zhang et al., 2015). Not only do we find that *Clstn3*, different from previously identified synaptic adhesion molecules, restricts formation at a specific synapse (parallel-fiber synapses), but also that *Clstn3* enhances formation of another specific synapse (GABAergic basket- and stellate-cell synapses) in the same neurons.

Several questions arise. First, why are the phenotypes we observe in *Clstn3* KO Purkinje cells so much stronger than those previously detected in CA1-region pyramidal neurons (Kim et al., 2020; Pettem et al., 2013)? This difference could be due to differences in cell type or to the more acute nature of our manipulations. More likely, however, this difference is caused by the lack of redundancy of *Clstn3* function in Purkinje cells since other calsyntenin isoforms are co-expressed with *Clstn3* in CA1-region neurons (Figure 1A), but not in Purkinje cells (Figure 1B, 1C).

Second, what is the mechanism of *Clstn3* action at synapses? We used manipulations in young adult mice in which cerebellar synapses are not yet completely established and in which synapse formation and elimination likely occurs continuously (Attardo et al., 2015; Pfeiffer et al., 2018). At present, our data do not reveal whether *Clstn3* acts in the initial establishment and/or the maintenance of synapses, a somewhat artificial distinction since synapse formation may actually consist in the stabilization of promiscuous contacts and synapses turn over continuously (Südhof, 2021). The functional consequences of these actions for cerebellar circuits are identical, in that both lead to a dramatic shift in excitatory/inhibitory balance in the cerebellar cortex.

Third, what trans-synaptic interactions mediate the functions of *Clstn3*? Several papers describe binding of calsyntenins to neurexins (Kim et al., 2020; Pettem et al., 2013). However, our data uncover a phenotype that is different from that observed with deletions of neurexins or neurexin ligands, suggesting that *Clstn3* does not function exclusively by binding to neurexins. The deletion of the neurexin ligand Cbln1 leads to a loss of parallel-fiber synapses in the cerebellar cortex instead of a gain, suggesting that a different calsyntenin ligand is involved. Moreover, the specific conclusions of the papers describing calsyntenin-binding to neurexins differ (Kim et al., 2020; Pettem et al., 2013), leaving the interaction mode undefined. Thus, we believe the most parsimonious hypothesis is that postsynaptic calsyntenins function by binding to presynaptic ligands other than neurexins that remain to be identified.

Fourth, does *Clstn3* physiologically act to restrict the formation of excitatory parallel-fiber synapses, leading to an increase in parallel-fiber synapses upon deletion of *Clstn3*, or is this increase an indirect compensatory effect produced by the decrease in inhibitory synapses? Multiple arguments support a specific action of *Clstn3* at parallel-fiber synapses. *Clstn3* protein was localized to parallel-fiber synapses by immunoelectron microscopy (Hintsch et al., 2002). Moreover, other genetic manipulations that cause a decrease in inhibitory synaptic transmission in cerebellar cortex, such as deletions of *Nlgn2* or of GABA_A_-receptors (Briatore et al., 2020; Fritschy et al., 2006; Meng et al., 2019; Zhang et al., 2015), do not induce an increase in excitatory parallel-fiber synapses. Finally and probably most importantly, although competition between synapses using the same transmitters is well-described (e.g., competition between glutamatergic parallel- and climbing-fiber synapses on Purkinje cells; Cesa and Strata, 2009; Miyazaki et al., 2012; Strata et al., 1997), no such competition has been observed between GABAergic and glutamatergic synapses, such that the decrease in one of them would lead to the increase of the other. Quite the contrary, the rules of homeostatic plasticity would predict that a decrease in GABAergic synapses should lead to a decrease, not an increase, in glutamatergic synapses (Monday et al., 2018; Nelson and Valakh, 2015). Thus, our data overall suggest that *Clstn3* specifically acts to limit the formation of parallel-fiber synapses and enhance the formation of inhibitory synapses in the cerebellar cortex.

Our study also has clear limitations. We did not examine axonal or dendritic transport, and cannot exclude the possibility that *Clstn3* performs an additional function as an adaptor for kinesin-mediated transport. Moreover, we cannot rule out the possibility that different calsyntenins perform distinct functions. In addition, although the postsynaptic functions of *Clstn3* in Purkinje cell synapses strongly argue against a neurexin-dependent mechanism, our data do not exclude the possibility that calsyntenins perform other functions in other neurons in a neurexin-dependent manner. The example of neuroligins shows that a synaptic adhesion molecule can have both a neurexin-dependent and neurexin-independent functions (Ko et al., 2009; Wu et al., 2019).

Multiple synaptic adhesion molecules have already been implicated in synapse formation in Purkinje cells. The interaction of presynaptic neurexins with cerebellins and postsynaptic GluD receptors plays a major role in shaping parallel-fiber synapses (Yuzaki and Aricescu, 2017), and the binding of C1ql1 to postsynaptic Bai3 has a prominent function in climbing-fiber synapses (Kakegawa et al., 2015; Sigoillot et al., 2015). Postsynaptic *Nlgn2* and *Nlgn3* are major contributors to the function of GABAergic synapses in Purkinje cells (Zhang et al., 2015), as is dystroglycan (Briatore et al., 2020). How can we envision the collaboration of various synaptic adhesion complexes in establishing and shaping the different types of synapses on Purkinje cells? Do these molecules act sequentially at different stages, collaborate, or work in parallel? The overall view of synapse formation that emerges from these studies resembles a conductorless orchestra, in which different players individually contribute distinct essential facets to the work that is being performed. In this orchestra, some players, such as neurexins, play prominent roles in coordinating the actions of their sections, whereas others, such as latrophilins, initiate movements. In this scenario, *Clstn3* (and possibly other calsyntenins) may regulate the loudness of different sections of the orchestra, or translated into the terms of a synapse, control the efficacy of signals regulating excitatory vs. inhibitory synapses.

## METHODS

### Key resources table

| Reagent type (species) or resource | Designation | Source or reference | Identifiers | Additional information |
| --- | --- | --- | --- | --- |
| Genetic reagent (Mus musculus) | Constitutive Cas9 | PMID: 25263330 | JAX ID: 024858 |  |
| Cell line (Homo sapiens) | HEK293T | ATCC | CRL-11268 |  |
| Recombinant DNA reagent | AAV-U6-sg66-U6-sg21-CAG tdTomato | This paper |  | Serotyped with AAV-DJ |
| Sequencebased reagent | Clstn1 RNA FISH probe | Advanced Cell Diagnostics | Cat: 542611 |  |
| Sequencebased reagent | Clstn2 RNA FISH probe | Advanced Cell Diagnostics | Cat: 542621 |  |
| Sequence based reagent | Clstn3 RNA FISH probe | Advanced Cell Diagnostics | Cat: 542631 |  |
| Sequence based reagent | Clstn3 qPCR primers and probe | This paper |  |  |
| Antibody | Anti-pan Clstn1/2 rabbit polyclonal | PMID: 24613359 |  | 1:1000 IB |
| Antibody | Clstn3 rabbit polyclonal | PMID: 24613359 |  | 1:1000 IB |
| Antibody | Anti-ACTB mouse monoclonal | Sigma | #A1978 | 1:10000 IB |
| Antibody | IRDye® 680LT Donkey anti-Rabbit | LI-COR | 926-68023 | 1:10000 IB |
| Antibody | IRDye® 800CW donkey anti-mouse | LI-COR | 926-32212 | 1:10000 IB |
| Antibody | Anti-vGluT1<br>rabbit polyclonal | Yenzym | YZ6089 | 1:1000 IHC |
| Antibody | Anti-vGluT2<br>rabbit polyclonal | Yenzym | YZ6097 | 1:1000 IHC |
| Antibody | Anti-vGAT<br>guinea pig polyclonal | Sysy | 131004 | 1:1000 IHC |
| Antibody | Alexa goat anti guinea pig<br>633 | Invitrogen | A-21105 | 1:1000 IHC |
| Antibody | Alexa goat anti rabbit 647 | Invitrogen | A-21245 | 1:1000 IHC |
| Chemical<br>compound, drug | Tribromoethanol | Sigma | T48402 | 250 mg/kg<br>for<br>anesthesia |
| Chemical<br>compound, drug | Picrotoxin | Tocris | 1128 |  |
| Chemical<br>compound, drug | APV | Tocris | 0106 |  |
| Chemical<br>compound, drug | CNQX | Tocris | 1045 |  |
| Chemical<br>compound, drug | NBQX | Tocris | 1044 |  |
| Chemical<br>compound, drug | Tetrodotoxin | Cayman<br>Chemical | 14964 |  |
| Chemical<br>compound, drug | DAPI | Sigma | D8417 |  |
| Chemical<br>compound, drug | Biocytin | Sigma | B4261 |  |
| Software,<br>algorithm | SnapGene | GSL<br>Biotech |  | previously<br>existing |
| Software,<br>algorithm | Image Studio Lite | LI-COR |  | previously<br>existing |
| Software,<br>algorithm | pClamp10 | Molecular<br>Device |  | previously<br>existing |
| Software,<br>algorithm | Clampfit10 | Molecular<br>Device |  | previously<br>existing |
| Software,<br>algorithm | NIS-Elements<br>AR | Nikon |  | previously<br>existing |
| Software,<br>algorithm | ImageJ | National<br>Institutes of<br>Health |  | previously<br>existing |
| Software,<br>algorithm | Neurolucida360 | MBF<br>science |  | previously<br>existing |
| Software, algorithm | Adobe Illustrator | Adobe |  | previously existing |
| Software, algorithm | Graphpad Prism 8.0 | Graphpad software |  | previously existing |

IB: immunoblotting, IHC: immunohistochemistry

### Animals

Constitutive Cas9 mice (https://www.jax.org/strain/024858) were used and maintained as homozygotes (Platt et al., 2014). Analyses were performed on littermate mice. Mice were fed ad libitum and on 12 hour light dark cycles. All protocols were carried out under National Institutes of Health *Guidelines for the Care and Use of Laboratory Animals* and were approved by the Administrative Panel on Laboratory Animal Care at Stanford University.

### Single molecule RNA fluorescent in-situ hybridization (smRNA-FISH)

smRNA-FISH in-situ hybridization experiment was performed on brain sections from P30 wild type C57BL/6J mice according to the manufacturer instructions using Multiplex Fluorescent Detection Reagents V2 kit (# 323110, Advanced Cell Diagnostics). Predesigned probes for Clstn1 (# 542611), Clstn2 (# 542621), and Clstn3 (# 542631) were purchased from ACD.

### SgRNA design and generation of Vectors

SgRNAs were designed using protocols developed by the Zhang lab (https://zlab.bio/guide-design-resources) to minimize potential off-target effects. The pAAV construct was modified from Addgene #60231 (Platt et al., 2014) with Cre-GFP replaced by tdTomato and human synapsin by CAG promoter to allow efficient expression in the cerebellum. Two sgRNAs were cloned in a single vector using Golden Gate Cloning assembly. Empty vector without sgRNAs was used as control. Genome editing efficiency of sgRNAs was initially evaluated using TIDE (https://tide.nki.nl/) (Brinkman et al., 2014).

Potential off-target editing sites were chosen from predictions while design. Forward and reverse primers were designed to flank sgRNAs, and PCR product of Genomic DNA were sequenced and compared on TIDE.

Primers for off-target site sequencing on sg66:

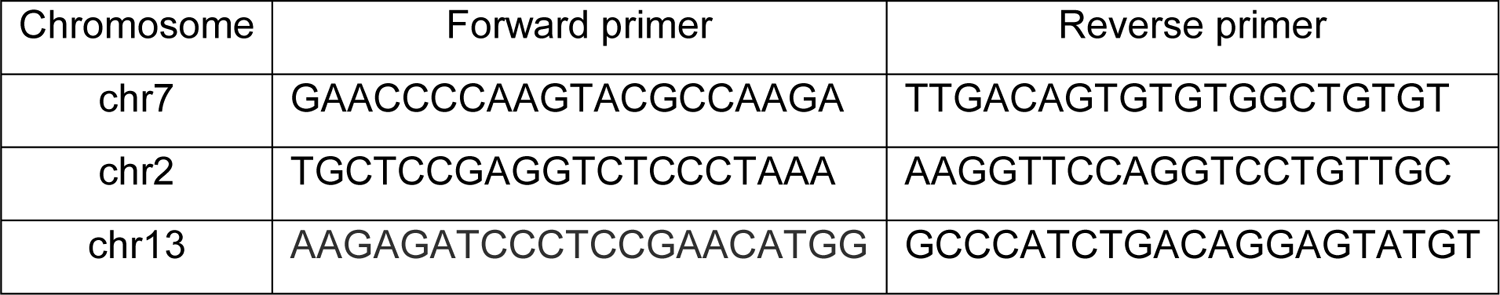

Primers for off-target site sequencing on sg21:

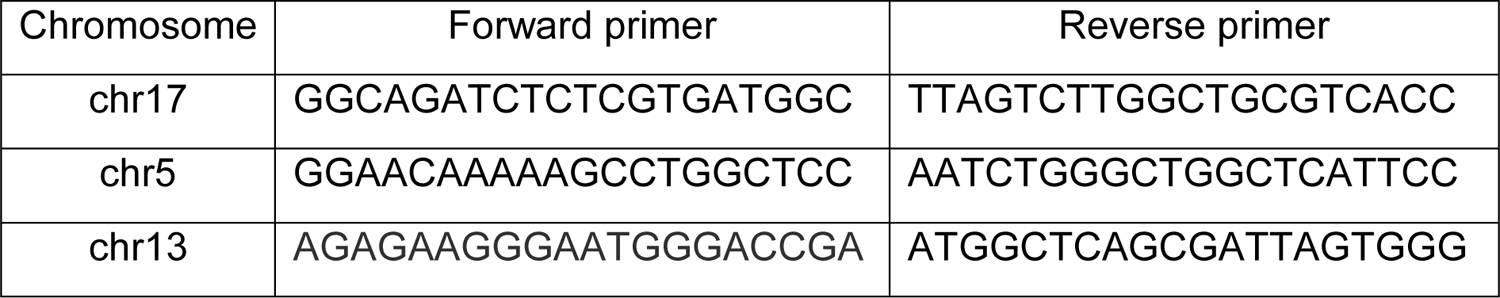

### AAV preparation and stereotactic Injections

pAAV carrying sgRNAs was serotyped with the AAV-DJ capsid (Grimm et al., 2008). Briefly, helper plasmids (phelper and pDJ) and AAV-sgRNA vector were co-transfected into HEK293T cells (ATCC, CRL-11268), at 4 μg of each plasmid per 30 cm^2^ culture area, using the calcium phosphate method. Cells were harvested 72 h post-transfection, and nuclei were lysed and AAVs were extracted using a discontinuous iodixanol gradient media at 65,000 rpm for 3 h. AAVs were then washed and dialyzed in DMEM and stored at −80 ℃ until use. Genomic titer was tested with qPCR and adjusted to 5 x 10^12^ particles/ml for *in vivo* injections.

P21 Cas9 mice were anesthetized with tribromoethanol (250 mg/kg, T48402, Sigma, USA), head-fixed with a stereotaxic device (KOPF model 1900). AAVs carrying sgRNAs or control viruses were loaded via a glass pipette connected with a 10 μl Hamilton syringe (Hamilton, 80308, US) on a syringe injection pump (WPI, SP101I, US) and injected at a speed of 0.15 μl/min. Pipette was left in cerebellum for additional 5 min after injection completion. Carprofen (5 mg/kg) was injected subcutaneously as anti-analgesic treatment. To infect the whole cerebellum, we injected multiple sites evenly distributing over the cerebellum skull, coordinates were as previously reported (Zhou et al., 2020), anterior to bregma, lateral to midline, ventral to dura (mm): (–5.8,±0.75), (–5.8, ±2.25), (–6.35, 0), (–6.35, ±1.5), (–6.35, ±3), (–7, ±0.75), and (–7, ±2.25), with a series of depth (mm): 2, 1.5, 1, 0.5, and volume was 0.3 μl/depth. Viruses were coded during virus injection and remained blinded throughout the whole study until data analyses were done.

### Quantitative RT-PCR

Virus-infected cerebellar tissue indicated by tdTomato was carefully dissected under fluorescence microscope. RNA was extracted using Qiagen RNeasy Plus Mini Kit with the manufacturer’s protocol (Qiagen, Hilden, Germany). Quantitative RT-PCR was run in QuantStudio 3 (Applied biosystems, Thermo Fisher Scientific, USA) using TaqMan Fast Virus 1-Step Master Mix (PN4453800, Applied biosystems, Thermo Fisher Scientific, USA). PrimerTime primers and FAM-dye coupled detection probes were used for detecting Clstn3 mRNA level. To detect genome editing efficiency, qPCR primers and probe were targeting the two exons and designed to flank the double-strand breaks of the two sgRNAs (Yu et al., 2014). (Clstn3: Forward primer: AGAGTACCAGGGCATTGTCA; reverse primer: GATCACAGCCTCGAAGGGTA; probe: TGGATAAAGATGCTCCACTGCGCT). A commercially-available GAPDH probe was used as internal control (Cat: 4352932E, Applied Biosystems).

### Immunohistochemistry

Immunohistochemistry on the cerebellar cortex was done as previously reported (Zhang et al., 2015). Mice were anesthetized with isoflurane and sequentially perfused with phosphate buffered saline (PBS) and ice cold 4% paraformaldehyde (PFA). Brains were dissected and post-fixed in 4% PFA overnight, then cryoprotected in 30% sucrose in PBS for 24 h. 40 μm thick sagittal sections of cerebellum were collected using a Leica CM3050-S cryostat (Leica, Germany). Free floating brain sections were incubated with blocking buffer (5% goat serum, 0.3% Triton X-100) for 1 h at room temperature, then treated with primary antibodies diluted in blocking buffer overnight at 4 ℃ (anti-vGluT1, Rabbit, YZ6089, Yenzym, 1:1,000; anti-vGluT2, Rabbit, YZ6097, 1:1,000, Yenzym; anti-vGAT, guinea pig, 131004, Sysy,1:1,000). Sections were washed three times with PBS (15 min each), then treated with secondary antibodies (Alexa goat anti guinea pig 633, A-21105, Invitrogen, 1:1,000; or Alexa goat anti rabbit 647, A-21245, Invitrogen, 1:1,000) for 2 h at room temperature. After washing with PBS 4 times (15 min each), sections were stained with DAPI (D8417, Sigma) and mounted onto Superfrost Plus slides with mounting media. Confocal images were acquired with a Nikon confocal microscope (A1Rsi, Nikon, Japan) with 60x oil objective, at 1024 x 1024 pixels, with z-stack distance of 0.3 μm. All acquisition parameters were kept constant within the same day between control and Clstn3 KO groups. Images were taken from cerebellar lobules IV/V. Images were analyzed with Nikon analysis software. During analysis, we divided the cerebellar cortex into different layers to compare *Clstn3* KO effects. We defined 0-40% as superficial molecular layer and 40-80% as molecular deep layers, 80-100% as PCL, and we analyzed and labeled GCL separately in vGAT staining.

### Immunoblotting

Immunoblotting was performed as described previously (Zhang et al., 2015). Mice were anesthetized with isoflurane and decapitated on ice, with the cerebellum dissected out and homogenized in RIPA buffer (in mM: 50 Tris-HCl pH7.4, 150 NaCl, 1% Triton X-100, 0.1% SDS, 1 EDTA) with protease inhibitor cocktail (5056489001, Millipore Sigma) and kept on ice for 30 min. Samples were centrifuged at 14,000 rpm for 20 min at 4 ℃, supernatant were kept and stored in −80 ℃ until use. Proteins were loaded onto 4-20% MIDI Criterion TGX precast SDS-PAGE gels (5671094, Bio-Rad), and gels were blotted onto nitrocellulose membranes using the Trans-blot turbo transfer system (Bio-Rad).

Membranes were blocked in 5% milk diluted in PBS for 1 h at room temperature, then incubated overnight at 4 ℃ with primary antibodies diluted in 5% milk in TBST (0.1% Tween-20). Primary antibodies of anti-Clstn3 (Rabbit, 1:1,000) and anti-pan Clstn1 and 2 (Rabbit, 1:1,000) were previously described (Um et al., 2014). Antibody against beta-actin from Sigma (A1978, Mouse, 1:10,000) was used as a loading control.

Membranes were then washed with TBST and incubated with fluorescence labeled IRDye secondary antibodies (IRDye® 680LT Donkey anti-Rabbit, 926-68023, LI-COR, 1:10,000; IRDye® 800CW donkey anti-mouse, 926-68023, LI-COR, 1:10,000). Signals were detected with Odyssey CLx imaging systems (LI-COR) and data were analyzed with Image Studio 5.2 software. Total intensity values were normalized to actin prior to control.

### Electrophysiology

Cerebellar electrophysiology was carried out as described previously (Caillard et al., 2000; Foster and Regehr, 2004; Llano et al., 1991; Zhang et al., 2015). Briefly, the cerebellum was rapidly removed and transferred into continuously oxygenated ice cold cutting solutions (in mM: 125 NaCl, 2.5 KCl, 3 MgCl_2_, 0.1 CaCl_2_, 25 glucose, 1.25 NaH_2_PO_4_, 0.4 ascorbic acid, 3 myo-inositol, 2 Na-pyruvate, and 25 NaHCO_3_). 250 μm sagittal slices were cut using a vibratome (VT1200S, Leica, Germany) and recovered at room temperature for >1 h before recording. Oxygenated ACSF (in mM: 125 NaCl, 2.5 KCl, 1 MgCl_2_, 2 CaCl_2_, 25 glucose, 1.25 NaH_2_PO_4_, 0.4 ascorbic acid, 3 myo-inositol, 2 Na-pyruvate, and 25 NaHCO_3_) was perfused at 1 ml/min during recording. Whole cell recordings with Purkinje cells were from cerebellar lobules IV/V, with patch pipettes (2-3 MΩ) pulled from borosilicate pipettes (TW150-4, WPI, USA) using PC-10 puller (Narishige, Japan). The following internal solutions were used (in mM): (1) for EPSC, 140 CsMeSO_3_, 8 CsCl, 10 HEPES, 0.25 EGTA, 2 Mg-ATP, 0.2 Na-GTP (pH adjusted to 7.25 with CsOH); (2) for IPSC, 145 CsCl, 10 HEPES, 2 MgCl_2_, 0.5 EGTA, 2 Mg-ATP, 0.2 Na-GTP (pH adjusted to 7.25 with CsOH). Liquid junction was not corrected during all recordings. For all EPSC recordings, 50 μM picrotoxin (1128, Tocris) and 10 μM APV (0106, Tocris) were contained in ACSF, and (1) additionally 0.5 μM NBQX (1044, Tocris) were included for climbing-fiber EPSC recordings; (2) 1 μM TTX (Tetrodotoxin, 14964, Cayman Chemical) for mEPSC recordings. For all IPSC recordings, (1) 10 μM CNQX (1045, Tocris) and 10 μM APV were included in ACSF, and (2) 1 μM TTX were included in mIPSC recordings. A glass theta electrode (64-0801, Warner Instruments, USA, pulled with PC-10) filled with ACSF was used as stimulating electrode, and littermate control and Clstn3 knockout mice were analyzed meanwhile using the same stimulating electrode. For climbing fibers, the electrode was placed in the granule cell layer around Purkinje cells and identified by all-or-none response, together with paired-pulse depression at 50 ms inter-stimulus interval (Eccles et al., 1966; Zhang et al., 2015). For parallel fibers, electrodes were placed in distal molecular layer with the same distance for both control and Clstn3 knockout mice (∼200 μm from the recorded Purkinje cell) and identified by paired-pulse facilitation at 50 ms inter-stimulus interval (Zhang et al., 2015). For basket cell stimulations, electrodes were placed in the proximal molecular layer (within∼100 μm from the recorded Purkinje cell) and also identified with all-or-none response (Caillard et al., 2000; Zhang et al., 2015). Cells with >20% changes in series resistances were rejected for further analysis. All electrophysiological data were sampled with Digidata1440 (Molecular Device, USA) and analyzed with Clampfit10.4. Coefficient of variation was calculated according to previous report (Lisman et al., 2007).

### Biocytin labeling in Purkinje cells

2 mg/ml Biocytin (B4261, Sigma) was dissolved in Cs-methanesulfonate internal solution followed by whole-cell voltage clamp recordings in the Purkinje cells (Sando et al., 2019). Slices were fixed in 4% PFA/PBS solution overnight at 4℃. Slices were then washed 3 × 5 mins with PBS, permeabilized, and blocked in 5% goat serum, 0.5% Triton-X100 in PBS at room temperature for 1 h. Then slices were incubated in 1:1,000 diluted Streptavidin Fluor™ 647 conjugate (S21374, Invitrogen) at room temperature for 2 h in 5% goat serum in PBS, washed 5 × 5 min with PBS, and mounted onto Superfrost Plus slides for imaging. Image overviews were obtained with a Nikon confocal microscope (A1Rsi, Nikon, Japan) with a 60x oil objective, at 1024 x 1024 pixels, with z-stack distance of 2 μm. Dendritic tree 3D reconstructions were performed using Neurolucida360 software (MBF science, USA) in the Stanford Neuroscience Microscopy Service Center. Note that some somas could not be detected automatically and were manually labeled. Spine images were obtained with a ZESS LSM980 inverted confocal, Airyscan2 for fast super-resolution setup, equipped with an oil-immersion 63X objective. Z-stacks were collected at 0.2 μm intervals at 0.06 μm/pixel resolution with Airyscan2. Spine images were deconvolved using ZEN blue software (ZESS). Spine density and characteristics were analyzed with Neurolucida360 software (MBF science, USA) in Stanford Neuroscience Microscopy Service Center. Only last order dendrites were analyzed, with 5-8 dendrites per cell per layer.

### Behavior

#### Accelerating rotarod

Mice were placed on an accelerating rotarod (IITC Life Science). The rod accelerated from 4 to 40 r.p.m. in 5 min. Mice were tested 3 times per day with 1 hour interval and repeated for 3 days. Time stayed on the rod was recorded while the mouse fell off, or hanged on without climbing, or reached 5 min.

#### Three-chamber social interaction

Social interaction was evaluated in a three-chamber box. Mice were placed initially in the central chamber to allow 10 min habituation for all three chambers. For sociability session, a same sex- and age matched stranger mouse (stranger1) was placed inside an upside-down wire pencil cup in one of the side chambers. The other side had the same empty pencil cup. A test mouse was allowed 10 min to investigate the three chambers. For social novelty session, another stranger mouse (stranger2) was placed into the empty pencil cup and test mouse was allowed another 10 min to investigate between three chambers. The time mice spent in each chamber was recorded and analyzed using BIOBSERVE III tracking system.

### Data analysis

Experiments and data analyses were performed blindly by coding viruses. Unpaired t-test or one-way ANOVA or two-way ANOVA or repeat measures ANOVA were used to analyze slice physiology data or immunohistochemistry data or behavior data as indicated in figure legends. Kolmogorov-Smirnov test was used to analyze the cumulative curves of mEPSCs or mIPSCs. Significance was indicated as *p < 0.05, **p < 0.01, ***p < 0.001. Data are expressed as means ± SEM.

### AUTHOR CONTRIBUTIONS

Z.L. performed AAV preparation, stereotactic injections, qRT-PCR, immunoblotting, immunohistochemistry, biocytin morphological study, cerebellar electrophysiology and behavior; M.J. designed sgRNAs, made the sgRNA vector and tested sgRNAs genome editing efficiency; K.L.A. performed in-situ hybridization; J.K. provided antibodies against Clstn1/2 and Clstn3, and R.S.Z. performed additional electrophysiological recordings. Z.L., M.J., and K.L.A. analyzed the data. Z.L. and T.C.S. wrote the manuscript.

## ACKNOWLEDGEMENTS

We thank Drs Bo Zhang, Justin Howard Trotter, Karthik Raju, Mu Zhou, and Richard Sando for advice, discussions, and help in this project. This paper was supported by a grant from NIH (MH052804 to T.C.S.).

## SUPPLEMENTARY INFORMATION

**Figure S1:**
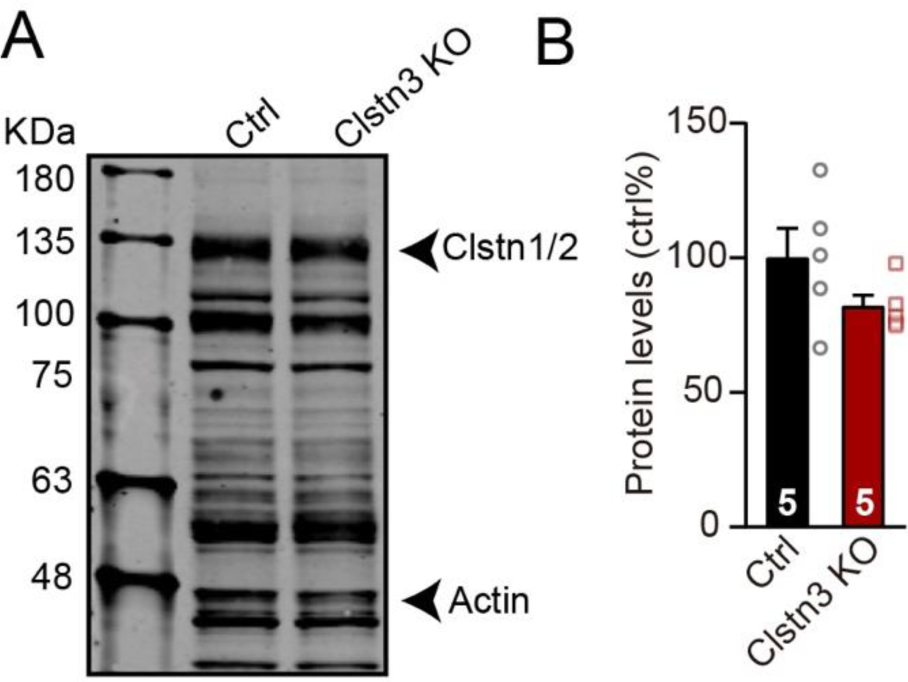
CRISPR-mediated deletion of Clstn3 does not cause a major change in the levels of Clstn1/2 protein in cerebellum. **(A)** Representative immunoblot of cerebellar homogenates from mice that were stereotactically injected in the cerebellum with Clstn3 CRISPR-KO or control AAVs. Note that the antibody used has innumerable non-specific crossreactivities, but that the bands identified as representing the combination of Clstn1 and Clstn2 (Clstn1/2) were previously validated for this antibody (Um et al., 2014). **(B)** Summary graph of the Clstn1/2 protein levels in control cerebellum and cerebellum with the Clstn3 KO. Data are means ± SEM (n = 5 mice). An unpaired t-test revealed no statistically significant difference between control and Clstn3 KO mice.

**Figure S2:**
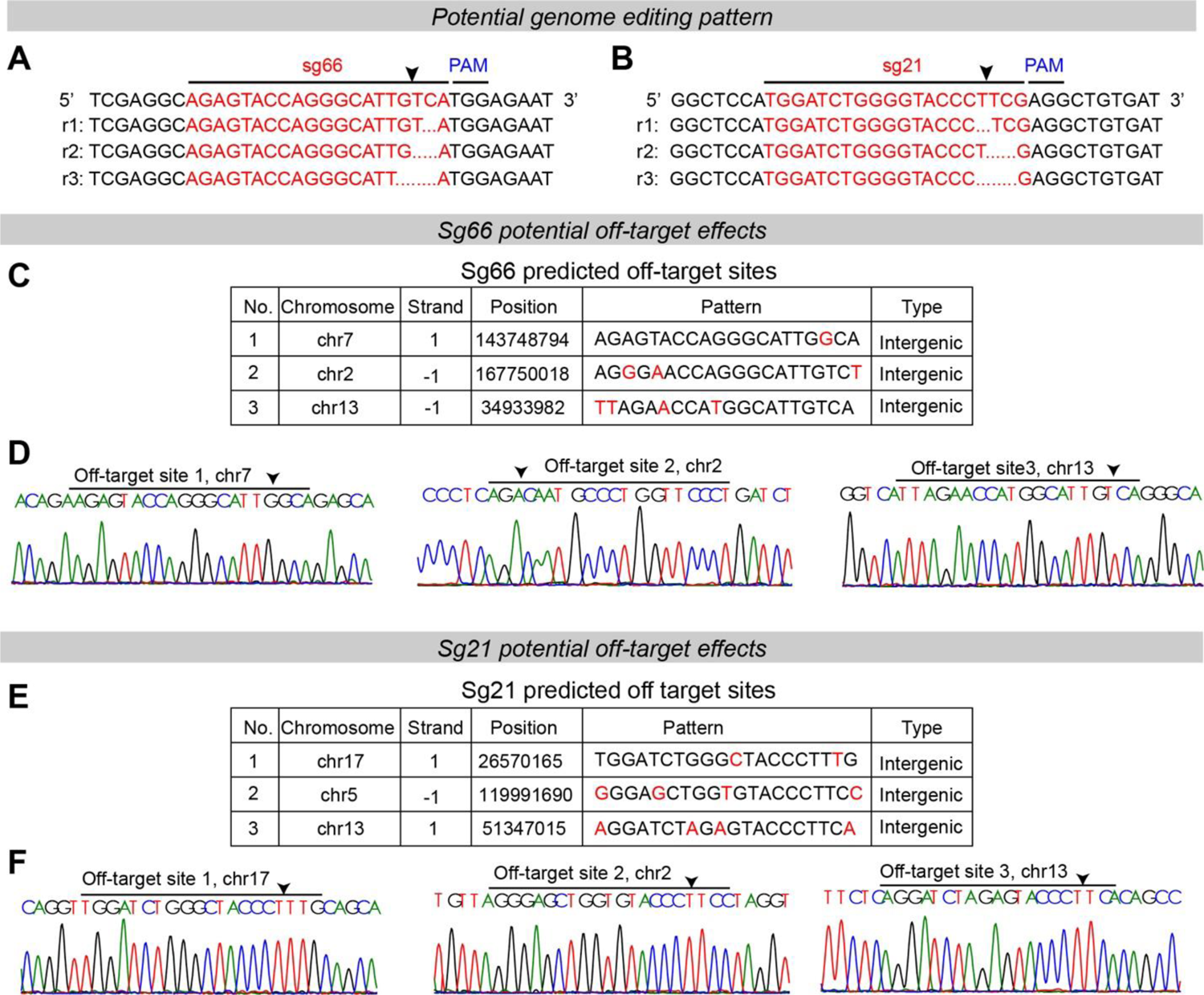
Predicted genome editing patterns by the sgRNAs used for the *Clstn3* KO in the current study and analysis of potential off-target effects of the sgRNAs using genomic sequencing of targeted *Clstn3* KO cerebellum. **(A-B)** Predicted genome editing effects for sg66 (A) and sg21 (B). **(C & D)** The three top-ranked potential off-target sites for sgRNA66 as analyzed by sequence predictions (C), and their analysis by genomic sequencing of *Clstn3* KO cerebellum, demonstrating no obvious gene editing effects on all three sites (D). Arrow indicates potential cutting positions. **(E & F)** Same as C & D, but for sgRNA21.

**Figure S3:**
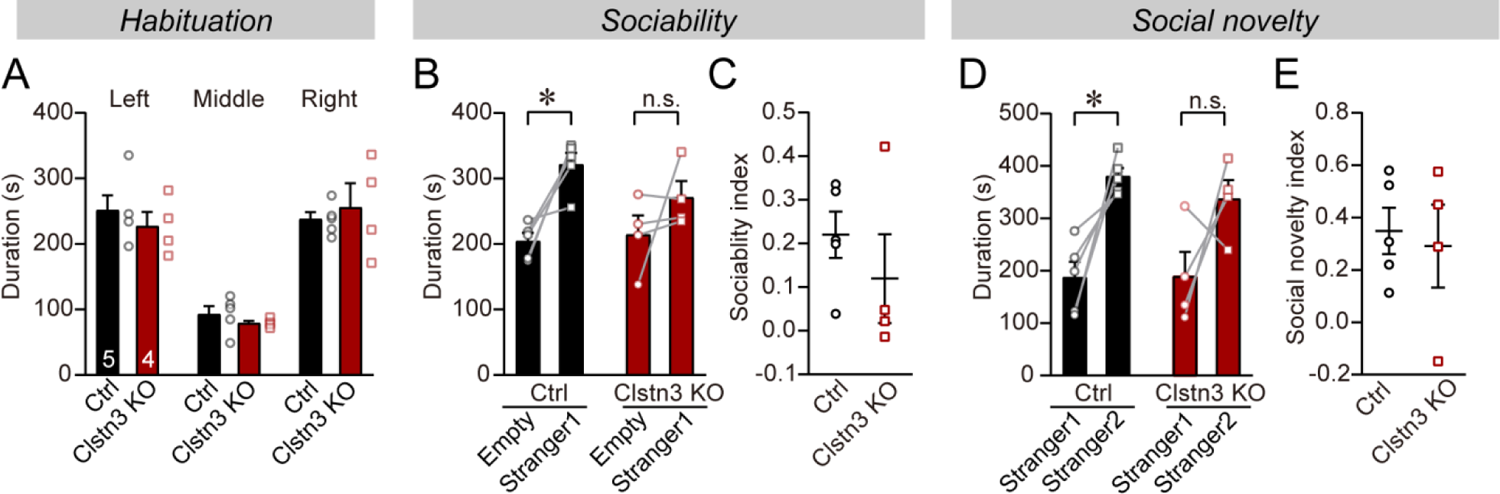
The *Clstn3* KO in the cerebellum does not significantly affect sociability and social novelty in mice as measured using the 3-chamber social behavior test. **(A)** Control and *Clstn3* KO mice exhibited the same exploration behavior of the left and right chambers during the habituation period. **(B & C**) When exposed to a non-familiar ‘stranger’ mouse in one of the outer chambers, cerebellar *Clstn3* KO mice exhibited a lower degree of interactions with the stranger mouse but this was not statistically significant (B, the time that the test mouse spent in the chambers with empty cup or ‘stranger1’ mouse; C, sociability index). **(D & E)** When given the choice between exploring a ‘stranger 1’ mouse to which it was previously exposed (see B), or a ‘stranger 2’ mouse that is novel, both control and cerebellar *Clstn3* KO mice prefer the novel mouse for interactions (D, the same as B, except empty cup has ‘stranger2’; E, social novelty index). All data are means ± SEM. Paired t-tests were applied to analyze the statistical significance of parameters within the same group, and unpaired t-tests to compare between control and Clstn3 KO groups at sociability index and social novelty index analysis, *p<0.05.

**Figure S4:**
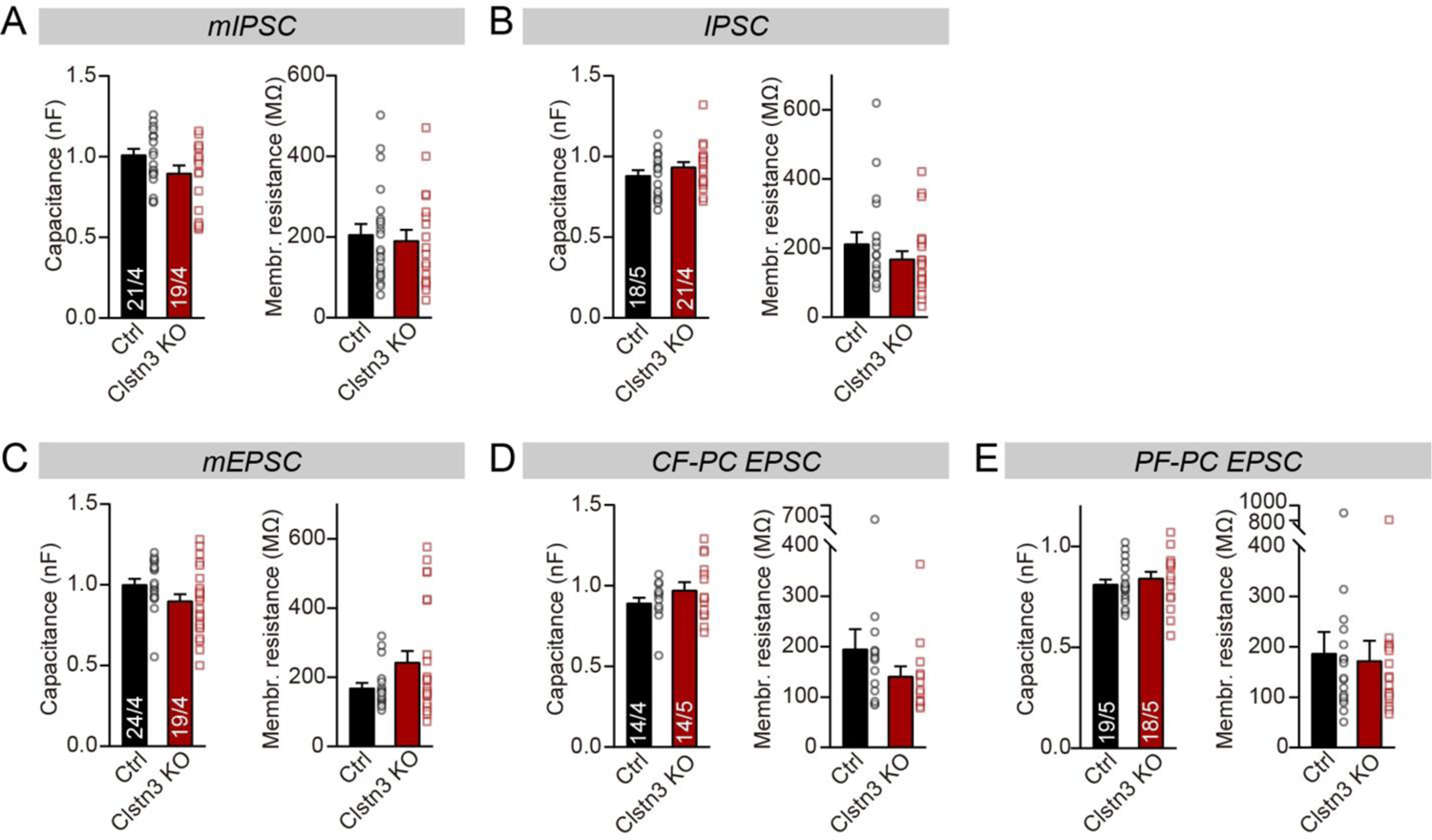
Capacitance and membrane resistance of Purkinje cells are unaffected by the *Clstn3* KO. **(A-E)** Summary graphs of the capacitance (left) and input resistance (right) of Purkinje cells determined during the recordings described in Fig. 5 and 6. All data are means ± SEM. Statistical analyses were done with unpaired t-test, p>0.05.

**Figure S5:**
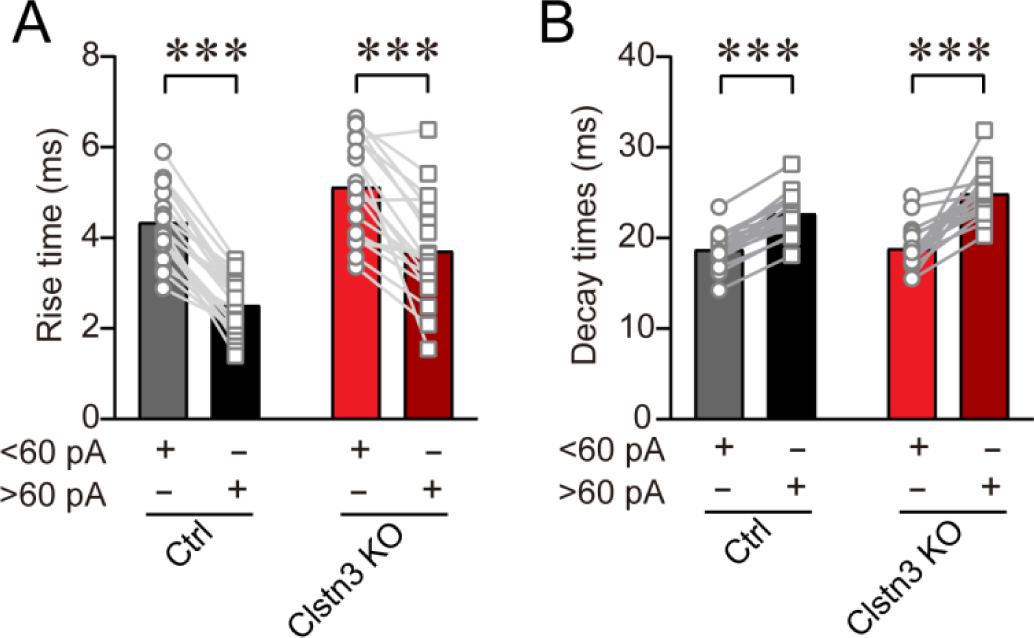
Analysis of the kinetics of large (>60 pA) and smaller (<60 pA) mIPSCs confirm that larger mIPSCs that are presumably generated by basket-cell synapses closer to the soma have faster rise times but slower decay times. **(A** & **B)** Rise (A) and decay times (B) of large and smaller mIPSCs examined in the same Purkinje cells. All data are means ± SEM. Statistical analyses were performed with paired t-test, ***p<0.001.

**Figure S6:**
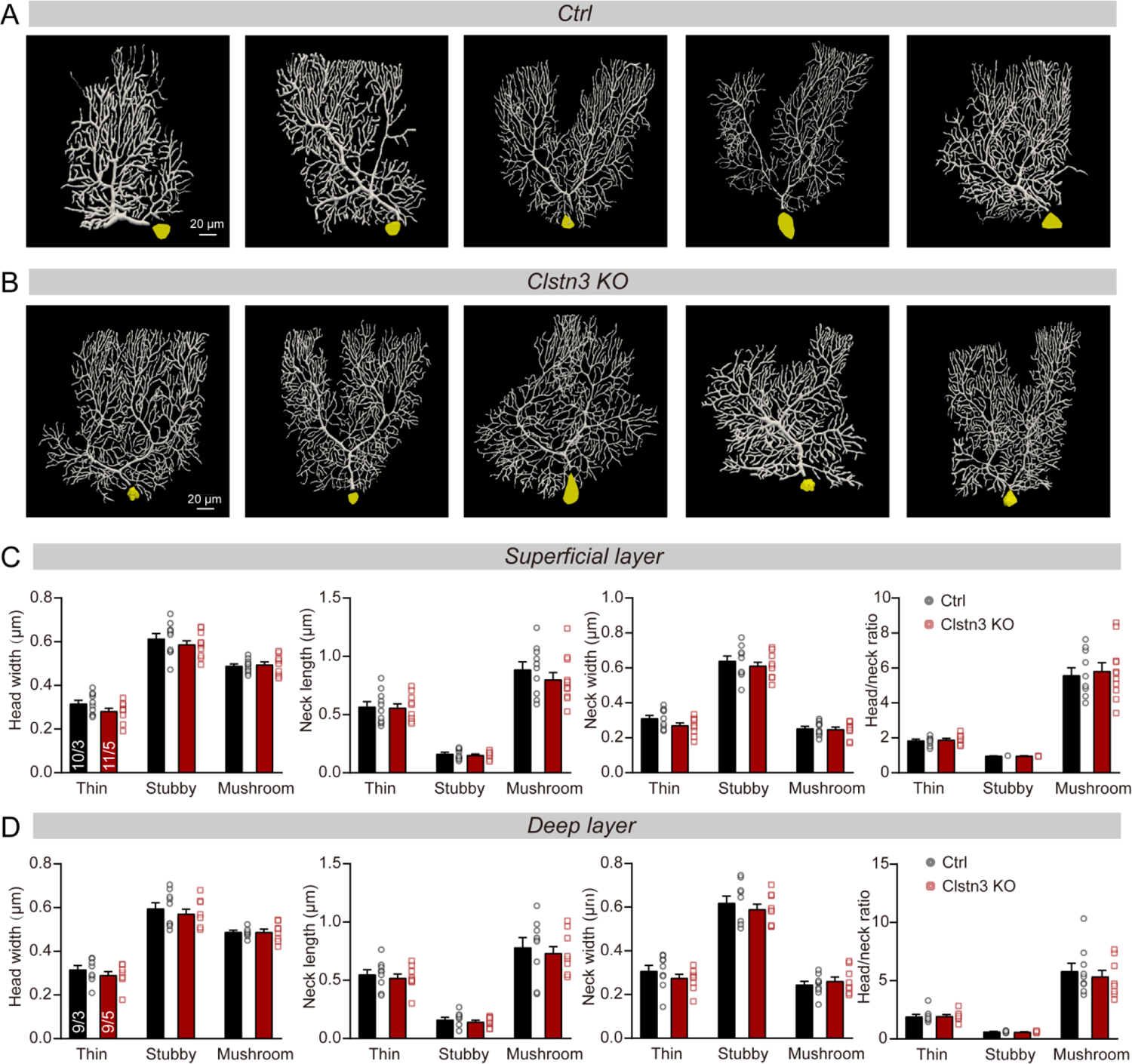
Images of individual reconstructed biocytin-filled Purkinje cells, and further quantifications of the morphological properties of spines from control and *Clstn3* KO Purkinje cells. **(A** & **B)** Images of all reconstructed Purkinje cells, as performed using Neurolucida360 software in control and *Clstn3* KO cerebellar cortex. (**C** & **D)** Quantification of various spine parameters in the superficial (C) and deep layers (D) of control and *Clstn3* KO mice (left, head width; middle left, neck length; middle right, neck width; right, head/neck ratio). Data are means ± SEM. 5-8 spines were analyzed per layer per cell; the numbers of cells/mice examined are indicated in the left bar graph. Statistical analyses were performed with unpaired Student’s t-tests, p>0.05.

**Figure S7:**
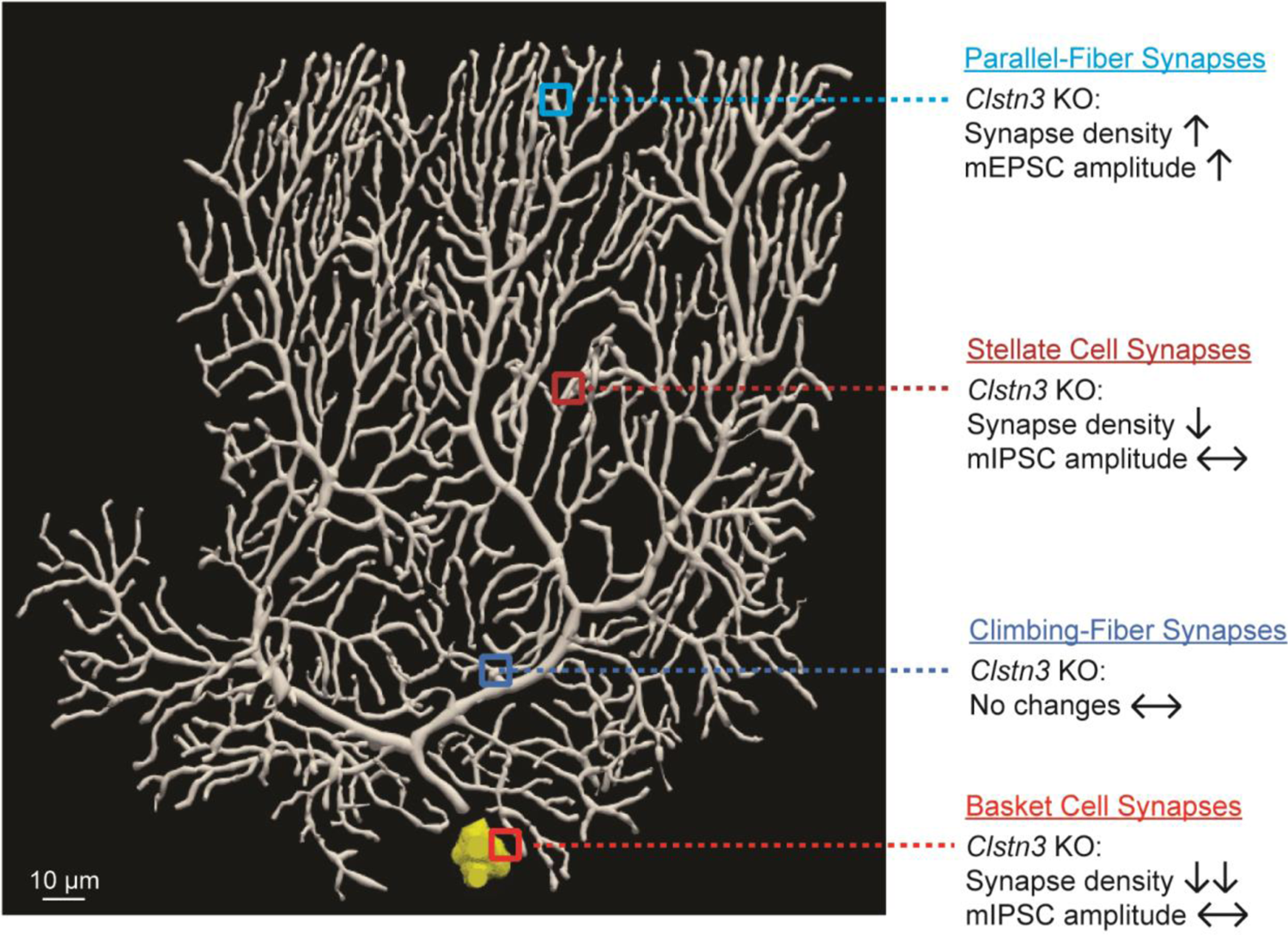
Summary of synaptic changes induced in Purkinje cells by the *Clstn3* KO. The Purkinje cell image is from one of the cells reconstructed during the present study. The changes summarized on the right were identified in Figures 3-9.

